# Neuronal late endosomes serve as selective RNA hubs disrupted by ALS-linked FUS mutation

**DOI:** 10.64898/2026.01.19.700337

**Authors:** Adriano Setti, Lorenzo Stufera Mecarelli, Tiziana Santini, Angelo D’Angelo, Vittorio Padovano, Erika Vitiello, Francesco Castagnetti, Julie Martone, Davide Mariani, Irene Bozzoni

## Abstract

Neurons depend on tightly regulated positioning of cellular components to maintain long-distance signaling, but the mechanisms guiding specific mRNAs to distant regions remain unclear. Here we show that late endosomes function as selective RNA carriers in human motor neurons and uncover the molecular logic guiding their loading. Using APEX2-mediated proximity labeling, we identify the external transcriptome of RAB7A-positive endosomes and find a specific population of mRNAs enriched for endosomal, axonal and synaptic functions. We find that mRNAs enriched in RAB7A endosomes contain evolutionarily conserved 5′UTRs which act as localization signals, and that the RNA-binding protein GEMIN5 interacts with these regions to promote endosomal RNA recruitment. Finally, we show that in ALS-associated conditions there is a conspicuous loss of endosome-associated transcripts and the mislocalization of GEMIN5 from endosomes. These findings uncover fundamental principles of RNA compartmentalization and highlight endosomal mRNA loading as a vulnerable axis in neuronal homeostasis.

## INTRODUCTION

The endosomal organelles are dynamic membrane-bound compartments that control the movement of cargos from the plasma membrane to lysosomes for degradation or back to the cell surface for recycling^1–3^. However, recent evidence has revealed that endosomes have a crucial role as mobile platforms for the transport of RNA–protein complexes in neurons. In particular, late endosomes can carry mRNAs on their cytosolic surface, enabling localized translation and targeted protein delivery within axons and dendrites^4–6^. Despite this emerging paradigm, the molecular mechanisms governing RNA recruitment to late endosomes remain poorly understood.

Disruptions in endosomal function and defects in axonal transport are well-established hallmarks of neurodegenerative diseases, including amyotrophic lateral sclerosis (ALS)^7^, where impaired vesicle trafficking contributes to neuronal dysfunction^8–10^. If RNAs fail to localize to endosomal carriers, local protein synthesis at distal sites such as synapses or mitochondria may be compromised, exacerbating neurodegeneration^11,12^.

One RNA-binding protein of particular interest in this context is FUS, a multifunctional RBP involved in transcription, splicing, RNA transport, and translation^13,14^. Importantly, FUS also participates in vesicle-mediated RNA trafficking^5^. Several ALS-causing mutations in FUS, such as P525L, determine its cytoplasmic mis-localization and aggregation^15^, disrupt the distribution of RBPs, such as SMN and FMRP, also involved in RNA transport^16,17^, and impair organelle mobility in axons^18^. Given its ability to bind both RNAs and RBPs and alter their subcellular localization^19^, mutant FUS could affect RNA loading onto late endosomes, with potential consequences for local translation and neuronal homeostasis. Understanding the composition of the RNA population associated with late endosomes, the intrinsic features that drive their localization, and how disease-associated mutations affect this process is therefore crucial.

In this study, we investigate the external transcriptome of late endosomes in motor neurons (MN), define some of the molecular features that promote RNA recruitment to these vesicles, and assess the impact of the ALS-linked FUS P525L mutation on the RNA composition of this compartment. To address these questions, we applied APEX2-mediated proximity labeling^20,21^ in human iPSC-derived motor neurons, enabling selective biotinylation and purification of RNAs residing on the cytosolic surface of late endosomes. This approach allowed us to generate the first comprehensive map of the late endosomal external transcriptome, revealing that neuronal late endosomes selectively associate mostly with mRNAs enriched in synaptic and endosomal functions, characterized by 5’UTRs bearing localization elements. We also identify GEMIN5 as one of the RBPs involved in targeting RNA to late endosomes. Strikingly, in FUS^P525L^ MN, we observed a dramatic loss of endosome-associated RNAs, a reduction in GEMIN5 recruitment to late endosomes, and an high binding affinity of mutant FUS for endosomal RNAs, indicating that this RBP disrupts endosomal RNA localization through both direct RNA binding and indirect interference with key RBPs.

Collectively, this work uncovers core principles governing RNA recruitment to late endosomes and reveals a previously unrecognized mechanism by which ALS-linked FUS mutations impair localized RNA transport, providing new insight into neurodegenerative disease pathogenesis.

## RESULTS

### The external transcriptome associated with RAB7A endosomes

The APEX2 proximity-labeling strategy enables biotinylation of proteins and RNAs within ∼20 nm from the fusion peptide after a brief hydrogen peroxide pulse^22^. Streptavidin pulldown then allows selective isolation and characterization of biotinylated molecules. We used this approach to isolate and sequence RNAs associated with the external surface of RAB7A-positive endosomes in human iPSC-derived MN (**Fig. 1A**).

**Figure 1.**
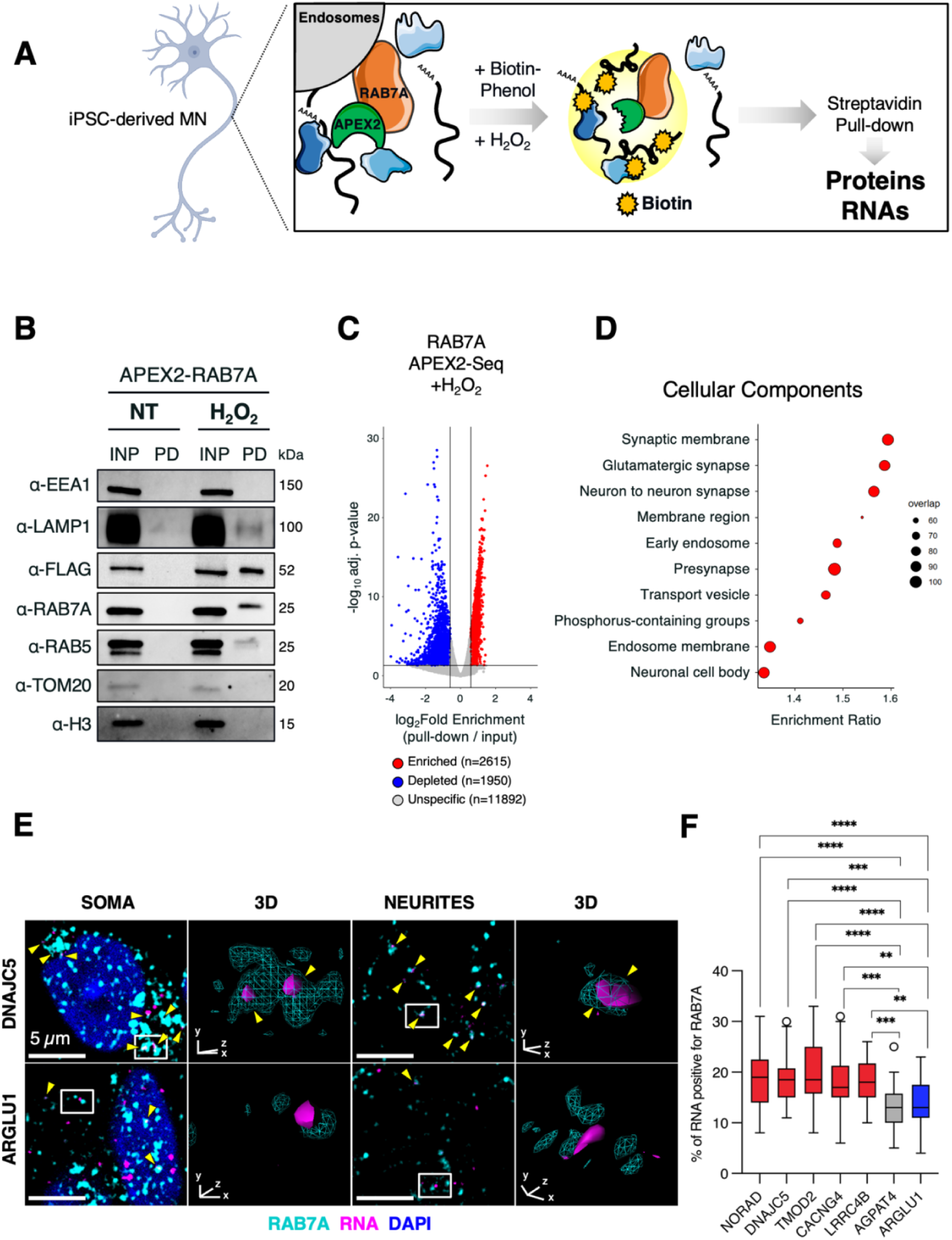
Proximity labeling reveals a distinct transcriptome associated with RAB7A-positive endosomes in motor neurons. **(A)** Schematic representation of the APEX2-based proximity labeling strategy used to biotinylate RNAs in the vicinity of RAB7A-positive endosomes in iPSCs-derived MNs. **(B)** Western blot analysis of the APEX2-pulldown on iPSCs-derived MNs expressing flagged APEX2-RAB7A protein. Early endosomal markers (EEA1, RAB5), Late endosomal marker (RAB7A), Late endosomal/lysosomal marker (LAMP1), APEX2-RAB7A construct (FLAG), Nuclear marker (H3), and mitochondrial marker (TOM20) were used to assess compartment purification. INP = 2,5% input, PD = pull down, H2O2= hydrogen peroxide treatment. **(C)** Volcano plot showing the log2FC pull-down/input and the −log10Pvalue for each RNA detected. Enriched and Depleted RNAs (Padj < 0.05, |log2FC| > 0.59) or Unspecific RNAs are indicated by red, blue and gray dots, respectively. **(D)** Dotplot depicting the cellular component GO over-represented categories of the *Enriched* RNAs. Only significant categories (FDR < 0.05) were depicted. X-axis represents category enrichment score, while Y-axis reports the GO category description. Dot size represents the amount of RNAs in the analyzed group that overlap the category. **(E)** Representative RNA FISH and immunofluorescence images showing localization of an endosome-*enriched* (DNAJC5) or endosome-*depleted* (ARGLU1) mRNA (magenta) relative to RAB7A-positive endosomes (cyan) in soma and neurites. Insets show 3D renderings of boxed regions. Scale bar = 5 μm. Colocalization is indicated by yellow arrows. **(F)** Quantification of the percentage of endosome-*enriched* (red), *unspecific* (gray), or *depleted* (blue) RNAs associated with RAB7A-positive particles. Data are shown as boxplots; p-values are indicated with asterisks (**p<0.01, ***p<0.001, ****p<0.0001). Statistical significance was assessed with One-way Brown-Forsythe and Welch ANOVA tests. N = 3 biological replicates (total cells: NORAD n = 315, DNAJC5 n = 270, TMOD2 n = 278, CACNG4 n = 272, LRRC4B n = 295, AGPAT4 n = 269, ARGLU1 n = 274).

iPSCs were engineered to stably express an APEX2-RAB7A construct with a N-terminal FLAG and subsequently differentiated into MN^23^. Immunofluorescence allowed us to confirm that the exogenous APEX2-RAB7A peptide co-localizes with the endogenous RAB7A positive foci (**Suppl. Fig. 1A**) and to prove that the overall number and distribution of endosomes in soma and neurites was comparable to control cells (**Suppl. Fig. 1B**). Moreover, Western blotting showed that exogenous APEX2-RAB7A is expressed at levels comparable to the endogenous RAB7A (**Suppl. Fig. 1C**). Analysis of the biotinylated proteins purified from APEX2-RAB7A MN confirmed the correct compartment targeting: the late endosomal constituent, RAB7A, was well represented, the late endosomal-lysosomal marker LAMP1 was detected at a minor extent, while the early endosomal proteins, EEA1 and RAB5, were at most barely detectable (**Fig. 1B**). As further controls, the signals corresponding to the nuclear histone H3 and to the mitochondrial TOM20 markers were absent (**Fig.1B**. Notably, none of these proteins were detected in cells not treated with hydrogen peroxide (“NT”, **Fig.1B**).

RNA-sequencing was performed by comparing pulldown versus the corresponding whole cell extracts. Among the 16457 RNAs expressed in MN (>1 FPKM in the input), we defined three groups: 2615 RNAs were enriched in the APEX2-RAB7A pull down fraction (defined as “*enriched*”; log₂ fold enrichment > 0.59, adjusted p < 0.05), 1950 species resulted depleted (“depleted”; log₂ fold enrichment < −0.59, adjusted p < 0.05), and 11892 were present at comparable levels in pull down and input samples ( “unspecific”, Fig. 1C). The extremely low number (5 species) of *enriched* RNAs detected in the untreated (NT) sample (**Suppl. Fig. 1D, Table S1**) demonstrated the specificity of the purification and the low background of the approach utilized.

The Gene Ontology (GO) analysis of the “*enriched*” RNAs showed the specific over-representation of categories related to synapses and endosomal membrane organization (**Fig. 1D, Suppl. Fig.1E and Table S1**). Moreover, KEGG pathway analysis further showed enrichment of genes involved in endocytosis and axonal development (**Suppl. Fig. 1F and Table S1)**. The relative abundance of RAB7A-positive particles in MN neurites, detected by immunofluorescence analysis (**Suppl. Fig. 1B**), supports the notion that several mRNAs encoding synaptic and endosomal proteins associate with endosomal membranes. This is consistent with the concept that endosomes act as platforms for the transport and localized translation of neurite-associated mRNAs^4^.

To validate the RNA-sequencing results, we performed in situ RNA FISH analysis to assess the co-localization of selected endosomal *enriched* transcripts (NORAD, DNAJC5, TMOD2, CACNG4, LRRC4B) with endogenous RAB7A. AGPAT4 and ARGLU1 mRNAs were used as *unspecific* and *depleted* controls, respectively (**Fig.1E**, **Fig.1F, Suppl. Fig. 1G** and **Suppl Fig. 1H**). Notably, all endosomal *enriched* RNAs showed significantly higher association with RAB7A-positive particles compared to the controls. Representative images of *enriched* (DNAJC5 and TMOD2) versus *depleted* and *unspecific* (ARGLU1 and AGPAT4) species are shown in **Fig.1E**, **Suppl. Fig. 1G**-**H**.

Collectively, these data demonstrate that late endosomes are enriched in MN neurites, and that their external surfaces are decorated mainly by RNAs encoding proteins involved in synaptic and endosomal functions, consistently with their role as hubs of mRNA localization and trafficking in neurons.

### 5’UTR sequences are sufficient to drive RNA localization to RAB7A endosomes

The data presented above demonstrate that a broad population of mRNAs is specifically associated with the external surface of late endosomes. This prompted us to investigate whether intrinsic features of these transcripts could mediate their selective recruitment to the appropriate subcellular compartment.

To identify RNA features driving transcript localization to endosomal membranes, we examined evolutionary conservation across 100 vertebrate species through phyloP scores and found that the 5′ UTRs of *enriched* RNAs are significantly more conserved than those of *depleted* transcripts (**Fig. 2A**). In contrast, the corresponding 3′ UTRs of both groups showed a lower degree of conservation (**Fig. 2A**), suggesting a possible role for the 5′UTR in targeting RNAs to endosomal membranes.

**Figure 2.**
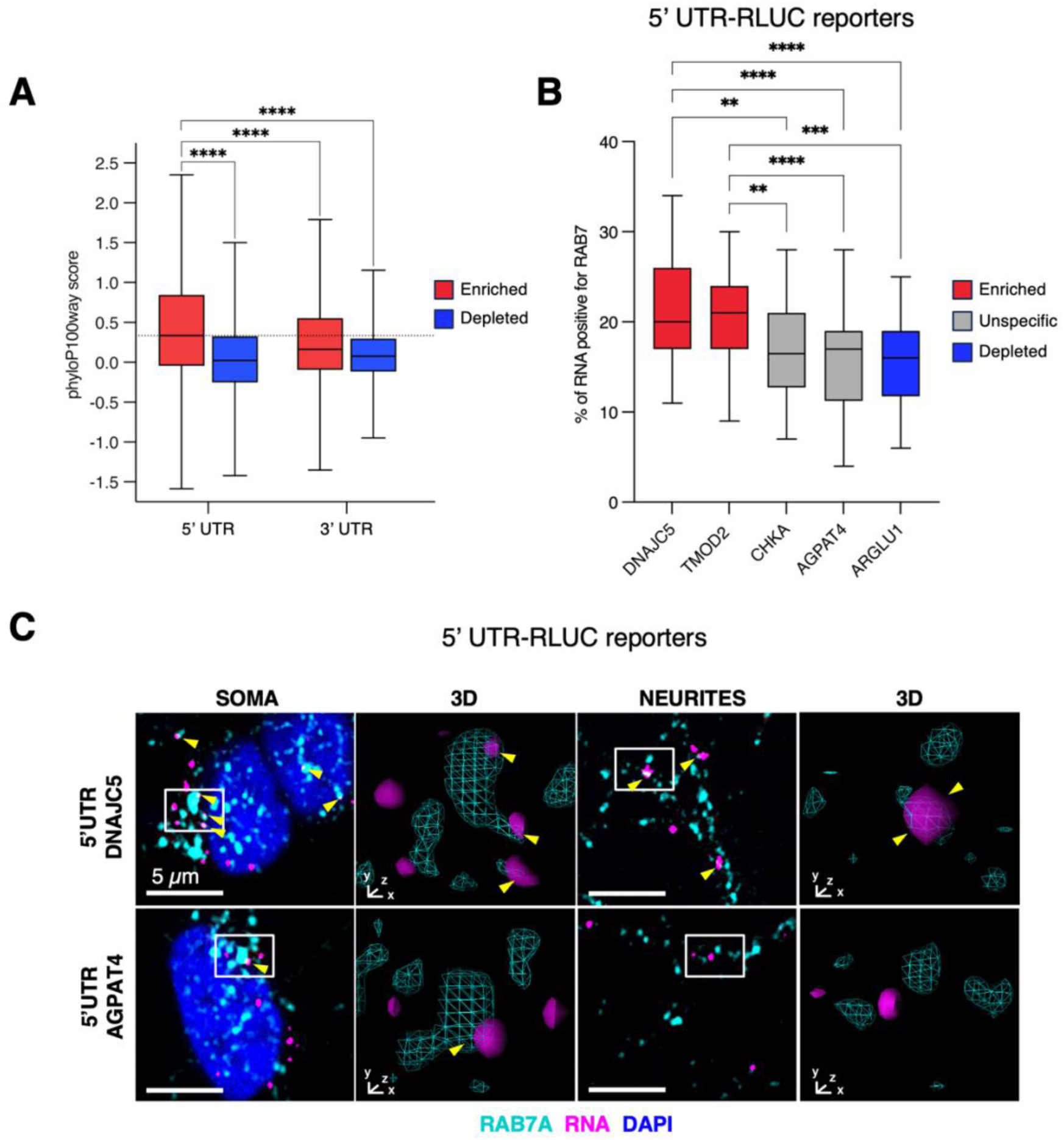
Conserved 5′UTR elements are sufficient to target RNAs to RAB7A-positive endosomes. **(A)** Distribution of PhyloP100 conservation scores for untranslated regions (UTRs). Boxplots show PhyloP100 scores for enriched (red) and depleted (blue) elements in the 5′ UTR (left pair) and 3′ UTR (right pair). **(B)** Quantification of the fraction of RLUC reporter mRNAs associated with RAB7A-positive endosomes when fused to 5′UTRs derived from endosome-*enriched* (red), *unspecific* (gray), or *depleted* (blue) transcripts. Data are shown as boxplots; p-values are indicated with asterisks (**p<0.01, ***p<0.001, ****p<0.0001). Statistical significance was assessed with One-way Brown-Forsythe and Welch ANOVA tests. N = 3 biological replicates (total cells: DNAJC5 n = 338, TMOD2 n = 354, CHKA n = 275, AGPAT4 n = 300, ARGLU1 n = 301). **(C)** Representative RNA FISH and immunofluorescence images showing localization of RLUC reporter mRNAs (magenta) fused to an endosome-*enriched* (DNAJC5) or *unspecific* (AGPAT4) 5′UTR relative to RAB7A-positive endosomes (cyan) in soma and neurites. 3D rendering of selected white boxes showing RAB7A-particles (cyan) and reporter mRNA (magenta). Scale bar = 5 μm. Colocalization is indicated by yellow arrows.

To test whether the 5’UTR regions of *enriched* RNAs are sufficient to mediate endosome association, we generated iPSC-derived MN stably expressing luciferase reporter constructs. The *Renilla* Luciferase ORF was fused to different 5′UTRs selected either among the endosome-*enriched* transcripts (DNAJC5 and TMOD2) or among different controls (CHKA*-unspecific*, AGPAT4-*unspecific* and ARGLU1*-depleted*). Co-localization between the reporter mRNAs and RAB7A was then assessed by immunofluorescence combined with RNA FISH. The data show that reporter RNAs carrying the 5′UTRs of endosome-*enriched* transcripts displayed significantly higher overlap with RAB7A-positive particles compared to those bearing control 5′ UTRs (**Fig. 2B**). Examples of FISH analysis for couples of *enriched* versus *depleted* RNAs with similar expression levels (**Suppl. Fig. 2A**) are shown in **Fig.2C** and **Suppl Fig. 2B**. Altogether, this evidence suggests that the 5’UTR of endosomal-*enriched* RNAs are sufficient to drive endosomal RNA localization.

### GEMIN5 binds the 5’UTR of endosome-associated mRNAs

RNA localization can be achieved through the binding of specific RBP, able to recognize binding sites usually located within the UTR sequences of RNAs^24–26^. In order to identify RBP possibly involved in the localization to endosomes, we analyzed the ENCODE eCLIP compendium^27^ focusing on the subgroup of those factors with predominant cytoplasmic localization (49 RBPs; see Methods). We identified four RBPs that preferentially bind the 5′ UTRs of endosome-*enriched* transcripts compared to *depleted* ones (FDR < 0.05, log2 OR > 0; **Fig. 3A** and **TableS2**). Notably, the four RBPs showed a specific binding within the 5′ UTRs of *enriched* transcripts, rather than in other regions (**Fig. 3B**).

**Figure 3.**
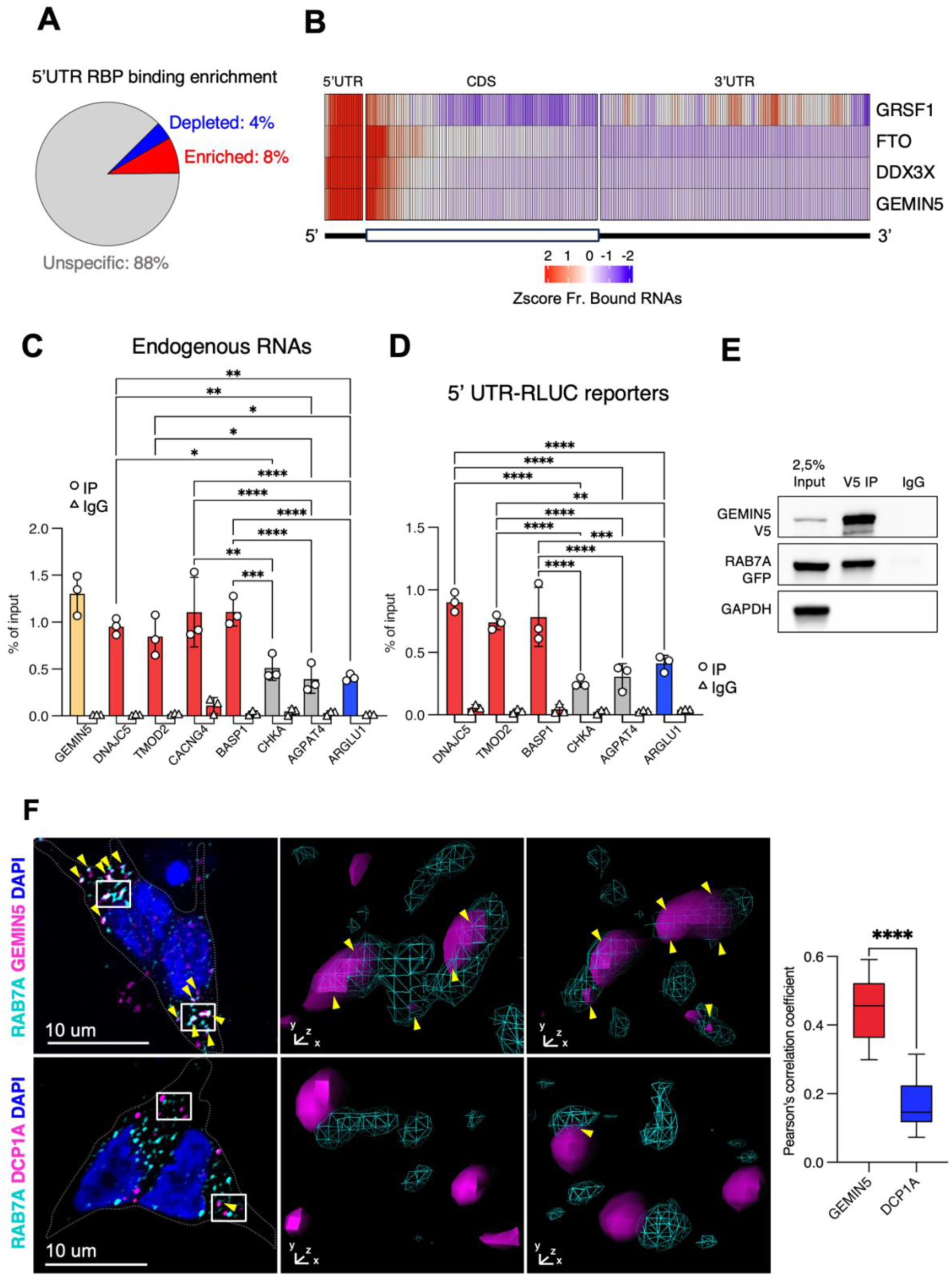
GEMIN5 binds the 5’UTR of endosome-*enriched* RNAs and localizes on LEs. **(A)** Pie chart showing the fraction of enriched, depleted or unspecific RBPs among 49 cytoplasmic RBPs from the ENCODE eCLIP compendium, as determined by RBP binding enrichment analysis on the 5′UTRs of RNAs enriched or depleted in endosomes **(B)** Meta-transcript heatmap showing the z-scored fraction of bound endosomal RNAs across transcript regions (5′UTR, CDS and 3′UTR) for the indicated RBPs (GRSF1, FTO, DDX3X and GEMIN5). Colour indicates relative enrichment (red) or depletion (blue) along the 5′–3′ axis. **(C)** qPCR quantification of GEMIN5 binding to endogenous endosome-*enriched* (red), *unspecific* (gray), and *depleted* (blue) RNAs in SK-N-BE cells, expressed as percentage of input. GEMIN5 mRNA serves as a positive control (yellow). p-values are indicated with asterisks (*p<0.05, **p<0.01, ***p<0.001, ****p<0.0001). Statistical significance was assessed with One-way ANOVA with Tukey multiple comparisons. **(D)** qPCR analysis of GEMIN5 binding to RLUC reporter mRNAs fused to 5′UTRs from endosome-*enriched* (red), *unspecific* (gray), or *depleted* (blue) transcripts in SK-N-BE cells, expressed as percentage of input. p-values are indicated with asterisks (*p<0.05, **p<0.01, ***p<0.001, ****p<0.0001). Statistical significance was assessed with One-way ANOVA with Tukey multiple comparisons. **(E)** Co-immunoprecipitation showing interaction between V5–GEMIN5 and GFP–RAB7A in neuroblastoma cells; GAPDH serves as a negative control. **(F)** Left, immunofluorescence analysis of GEMIN5 or the P-body marker DCP1A (magenta) relative to RAB7A-positive endosomes (cyan). Insets show 3D renderings of boxed regions. Scale bar = 10 μm. Right, quantification of colocalization by Pearson’s correlation coefficient. Data are shown as boxplots; p-value is indicated with asterisks (****p<0.0001). Statistical significance was assessed with two-tailed, unpaired T-test. N = 3 biological replicates (total cells: GEMIN5 n = 323, DCP1A = 259)

Among the four candidates—GEMIN5, FTO, DDX3X, and GRSF1—we selected GEMIN5 as the most plausible factor involved in recruiting and transporting RNA to endosomal membranes. Unlike the ubiquitously distributed demethylase FTO, the broadly expressed helicase DDX3X, or the mitochondria-enriched RBP GRSF1, GEMIN5 is known to form cytoplasmic RNA granules and to function as a selective mRNA-carrying RNP hub in neurons, exhibiting dynamics that closely mirror those of endosome-associated mRNAs^28^.

To assess whether GEMIN5 directly binds endosomal-*enriched* RNAs, we generated SK-N-BE neuroblastoma cell lines stably expressing a V5-tagged GEMIN5 protein. The immunoprecipitation of GEMIN5-V5 upon UV crosslinking revealed, with comparison to the positive control (GEMIN5) and to the negative (CHKA, AGPAT4 and ARGLU1) ones, a preferential binding of the protein to the endosomal *enriched* transcripts (DNAJC5, TMOD2, BASP1, CACNG4; **Fig. 3C** and **Suppl. Fig. 3B**). To further assess the specificity of GEMIN5 binding to the 5′ UTRs of endosome-enriched RNAs, we generated SK-N-BE cells co-expressing GEMIN5-V5 together with individual 5′UTR–RLUC reporter constructs. CLIP analysis in these cells (**Suppl. Fig. 3A**) demonstrated that GEMIN5 binds the 5′UTR–RLUC transcripts derived from endosome-enriched RNAs in a significantly higher manner with respect to control reporters (**Fig. 3D**). Notably, the efficiency of IP was comparable among the experiments (**Suppl. Fig. 3A, 3C and 3D**). Consistently, co-immunoprecipitation assays (**Fig. 3E**) and immunofluorescence in MN (**Fig. 3F),** confirmed the interaction between GEMIN5 and RAB7A. Moreover, GEMIN5 displays a higher and statistically significant level of co-localization with RAB7A than the negative control DCP1A (**Fig. 3F** and **Suppl. Fig. 3E**).

Taken together, these results indicate that GEMIN5 localizes to late endosomes, specifically binds endosomal-*enriched* RNAs through their 5′UTRs, and may act as a driver of RNA localization onto these vesicles.

### GEMIN5 localizes mRNAs to RAB7A endosomes through binding to their 5’UTR

To determine whether GEMIN5 plays a functional role in regulating endosomal RNA localization, we performed its downregulation through RNAi in SK-N-BE cells and analyzed the localization of its RNA interactors using immunofluorescence for RAB7A combined with RNA FISH (**Suppl. Fig. 4A**). We examined two endosome-*enriched* GEMIN5 RNA interactors (DNAJC5 and TMOD2; see **Fig. 1E and 3C**), together with two control RNAs that neither associate with endosomes nor interact with GEMIN5 (ARGLU1, *depleted*, and AGPAT4, *unspecific*; see **Fig. 1E and 3C**). Quantification of RNA spots colocalizing with RAB7A showed that GEMIN5 depletion significantly impaired the endosomal localization of *enriched* RNAs, whereas control transcripts remained unaffected (**Fig. 4A and Fig. 4B**). Consistently, the same experiment using the corresponding 5′UTR-RLUC reporters yielded similar results: DNAJC5 and TMOD2 constructs showed reduced endosomal colocalization upon GEMIN5 knockdown, while the AGPAT4 and ARGLU1 controls were unaffected (**Fig. 4B, Suppl. Fig 4B and 4C**). Taken together, these data demonstrate that GEMIN5 is required for proper endosomal localization of *enriched* RNAs that harbour a 5′UTR capable of interacting with this protein.

**Figure 4.**
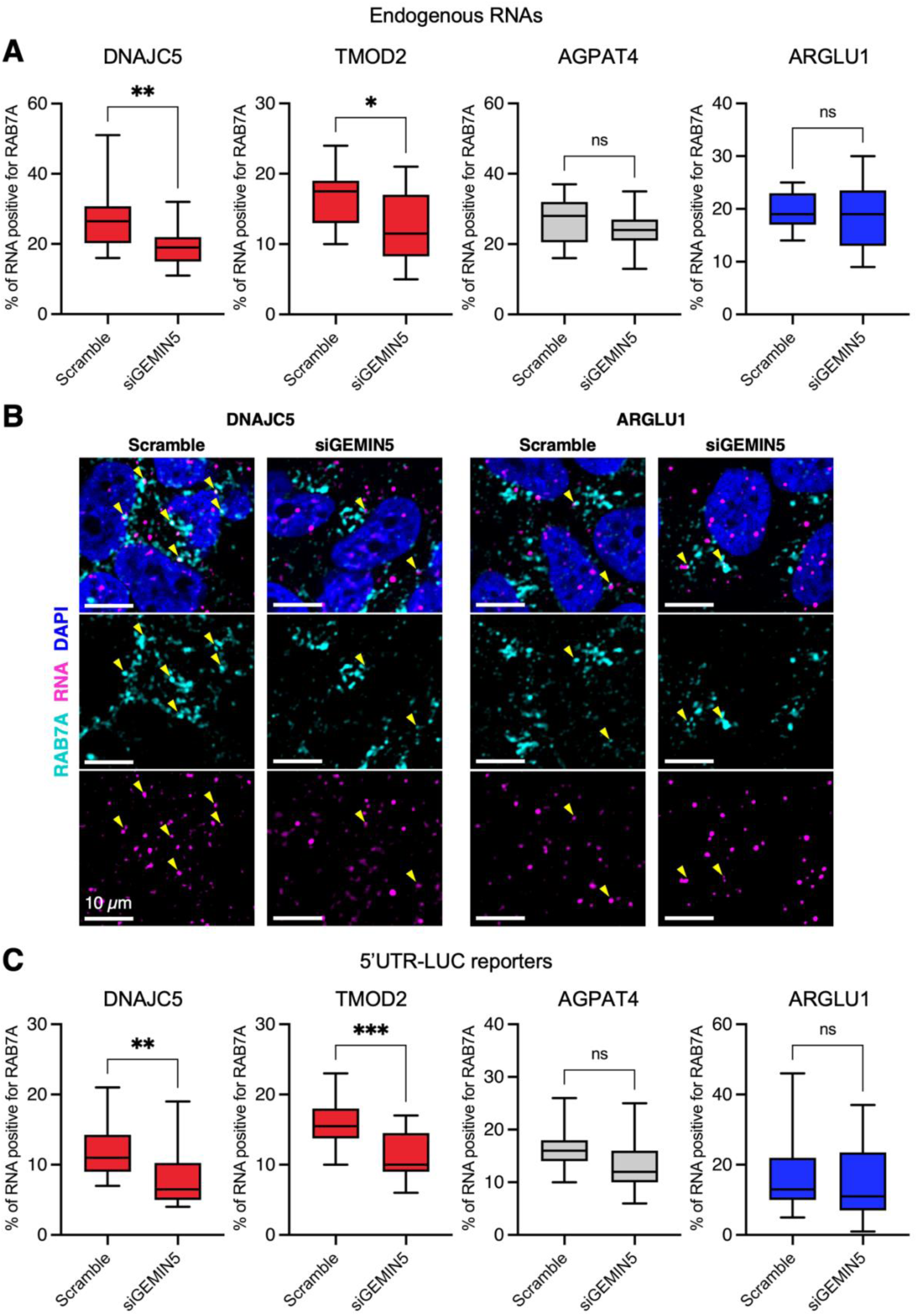
GEMIN5 is required for 5′UTR-dependent RNA recruitment to RAB7A-positive endosomes. **(A)** Quantification of endogenous RNA association with RAB7A-positive endosomes in control (Scramble) and GEMIN5 knockdown (siGEMIN5) SK-N-BE cells. Endosome-*enriched* (red), *unspecific* (gray), and *depleted* (blue) transcripts are shown. Data are shown as boxplots; p-values are indicated with asterisks (*p<0.05, **p<0.01). Statistical significance was assessed with two-tailed, unpaired T-test. **(B)** Representative RNA FISH and immunofluorescence images showing localization of an endosome-*enriched* (DNAJC5) or endosome-*depleted* (ARGLU1) mRNA (magenta) relative to RAB7A-positive endosomes (cyan) in control (Scramble) and GEMIN5 knockdown (siGEMIN5) SK-N-BE cells. Scale bar = 10 μm. Colocalization is indicated by yellow arrows. **(C)** Quantification of RAB7A association for RLUC reporter mRNAs fused to 5′UTRs from endosome-*enriched* (red), *unspecific* (gray), or *depleted* (blue) transcripts in control and GEMIN5 KD conditions. Data are shown as boxplots; p-values are indicated with asterisks (**p<0.01, ***p<0.001). Statistical significance was assessed with two-tailed, unpaired T-test.

### The ALS-associated mutant FUS^P525L^ impairs RNA recruitment to RAB7A endosomes

Recently, disruption of vesicular trafficking has clearly emerged as a hallmark of multiple neurodegenerative diseases, including ALS^29,30,9^. In particular, ALS-associated mutant FUS proteins, which mislocalize to the cytoplasm where they promote aggregate formation, have been shown to sequester key RBPs involved in RNA transport, such as IMP1^31^, SMN^16^ and kinesin1^32^. Therefore, we investigated the impact of the ALS-linked FUS^P525L^ mutation on the composition of the endosomal transcriptome.

To assess whether this mutation alters the endosomal transcriptome, we performed APEX2 - RAB7A-seq in isogenic iPSC-derived MN homozygous for FUS^P525L^. Efficient and specific biotinylation of endosomal proteins was confirmed by streptavidin-based pulldown followed by Western blotting (**Suppl. Fig. 5A**). Strikingly, RNA-seq analysis revealed a major collapse of the endosomal transcriptome. Of the 17,760 RNAs expressed (>1 FPKM in input), only 329 were enriched in FUS^P525L^ MN, compared to 2,615 in control cells (**Fig. 5A**). In addition, 662 were depleted, and 16,768 remained unchanged (**Suppl. Fig. 5B** and **Table S1**) with no *enriched* RNAs in the untreated (NT) samples (**Suppl. Fig. 5C**). Thus, approximately 91% of RNAs normally enriched at late endosomes in wild-type MNs were depleted in the FUS^P525L^ background, indicating a profound defect in endosomal RNA recruitment. To validate these findings, we performed immunofluorescence coupled with RNA FISH on transcripts that were selectively lost from endosomes in FUS^P525L^ conditions. As shown in **Fig. 5B** and **5C**, CACNG4 and TMOD2 RNAs, classified in the “*loss*” fraction by our sequencing analysis, exhibited a significant reduction in the association with RAB7A-positive endosomes. Representative examples for CACNG4 and TMOD2 mRNAs are shown in **Fig. 5C** and **Suppl. Fig. 5D**, respectively. By contrast, NORAD, belonging to the “*common*” group, and AGPAT4, excluded from endosomes in both conditions, maintained their localization even in FUS^P525L^ cells (**Fig. 5B**).

**Figure 5.**
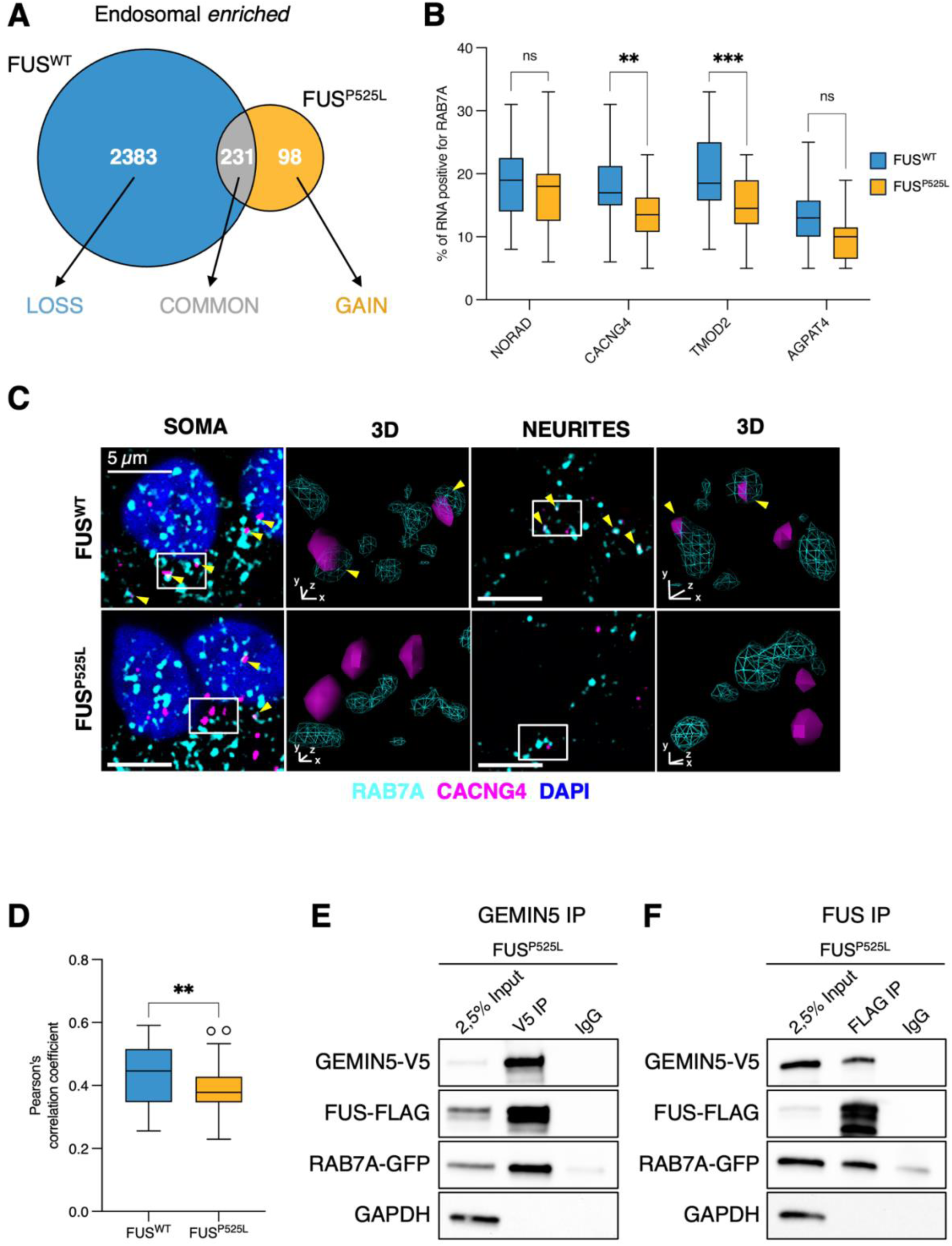
ALS-related FUS^P525L^ disrupts endosomal RNA recruitment. **(A)** Venn diagram showing the overlap between endosome-*enriched* RNAs identified in FUS^WT^ and FUS^P525L^ iPSCs-derived MNs. **(B)** Quantification of RAB7A association for representative RNAs that are commonly enriched (NORAD), selectively lost in FUS^P525L^ neurons (CACNG4, TMOD2), or commonly excluded (AGPAT4). Data are shown as boxplots; p-values are indicated with asterisks (*p<0.05, **p<0.01, ***p<0.001). Statistical significance was assessed with a two-way ANOVA test. N = 3 biological replicates (total cells: FUS^WT^ NORAD n = 315, CACNG4 n = 272, TMOD2 n = 278, AGPAT4 n = 269; FUS^P525L^ NORAD n = 349, CACNG4 n = 257, TMOD2 n = 282, AGPAT4 n = 260). **(C)** Representative RNA FISH and immunofluorescence images showing CACNG4 mRNA (magenta) relative to RAB7A-positive endosomes (cyan) in soma and neurites, in FUS^WT^ and FUS^P525L^ MNs. Scale bar = 5 μm. Colocalization is indicated by yellow arrows. 3D rendering of selected white boxes showing RAB7A-particles (cyan) and CACNG4 mRNA (magenta). **(D)** Pearson’s correlation coefficient analysis of GEMIN5 and RAB7A colocalization, in FUS^WT^ and FUS^P525L^ MNs. Data are shown as boxplots; p-values is indicated with asterisks (**p<0.01). Statistical significance was assessed with two-tailed, unpaired T-test. N = 3 biological replicates (total cells: FUS^WT^ n = 323, FUS^P525L^ = 298) **(E)** Co-IP for V5-GEMIN5 showing its interaction with FLAG-FUS^P525L^ and GFP-RAB7A in SK-N-BE cells. GAPDH serves as a negative control. **(F)** Co-IP for FLAG-FUS^P525L^ demonstrating its interaction with V5-GEMIN5 and GFP-RAB7A. GAPDH serves as a negative control.

Given the role of GEMIN5 in promoting endosomal RNA localization, we next asked whether its recruitment to late endosomes is impaired in the presence of mutant FUS^P525L^. Immunofluorescence analysis revealed a significant reduction in the colocalization between GEMIN5 and RAB7A, as quantified by Pearson’s correlation coefficient (**Fig. 5D**). Notably, GEMIN5 CLIP experiments showed that the interaction between GEMIN5 and CACNG4 was not affected in FUS^P525L^ cells (**Suppl. Fig. 5E**), indicating that the exclusion of CACNG4 RNA from endosomes is not due to a failure in GEMIN5 binding. Interestingly, we also found that FUS interacts with GEMIN5, as demonstrated by reciprocal co-immunoprecipitation (**Fig. 5E and 5F**). Altogether, these findings support a model in which mutant FUS sequesters GEMIN5, leading to defects in localization to endosomes for a specific class of transcripts.

Since FUS can also directly bind RNA, we tested whether it could subtract endosomal *enriched* RNA through direct binding. Therefore, we interrogated previously published PAR-CLIP data for FUS^P525L^ generated in human iPSC-derived MN^33^ to assess the binding propensity of mutant FUS toward endosomal RNAs. Interestingly, we found that FUS^P525L^ has high binding propensity for endosomal RNAs and that 48% of the 822 RNAs with the highest affinity for FUS (top 10% based on the proportion of PAR-CLIP transitions) are depleted from endosomes in the presence of mutant FUS (**Suppl. Fig. 5F**). Notably, immunofluorescence analysis revealed no significant alteration in RAB7A-positive particles number in FUS^P525L^ MNs (**Suppl. Fig. 5G**).

Taken together, these results reveal a profound collapse of the endosomal transcriptome in ALS-associated conditions. The concomitant displacement of GEMIN5, FUS, and their RNA targets from RAB7A-positive endosomes indicates that FUS^P525L^ disrupts RNA recruitment via both direct and indirect mechanisms.

## DISCUSSION

By defining the external transcriptome of RAB7A-positive late endosomes in iPSC-derived human MN, we reveal that these vesicles are associated with mRNAs encoding endosomal, axonal and synaptic regulators. These data, together with the notion that many of these transcripts are known to be localized at the neuronal periphery^34^, support a direct role for endosomes as platforms for mRNA trafficking and, possibly, local translation in neurites. This extends previous observations that endosomes transport mRNA–protein complexes in neurons and establishes a molecular framework for RNA recruitment to this compartment.

Mechanistically, we identify the 5′UTR of the mRNAs associated with APEX2-RAB7A endosomes as a key element governing RNA localization. These regions alone can direct mRNAs to late endosomes, and, for a specific subclass, we found that the RBP GEMIN5 mediates endosomal loading of selected mRNAs. While GEMIN5, being part of the SMN complex, has been implicated in RNA transport^28^ and neuronal ribonucleoprotein granule dynamics, our data position it as a previously unrecognized mediator linking endosomal vesicles to selective RNA cargo. This expands the functional repertoire of GEMIN5 beyond its canonical roles in snRNP assembly and axonal RNA granules, suggesting that neurons may utilize shared RNA-binding factors to coordinate distinct localization pathways converging on translation-competent organelles. Moreover, contrary to prevailing models that emphasize 3′UTRs as the primary drivers of mRNA localization, our data highlight a major role for the 5′UTRs in mediating mRNA targeting to endosomal membranes. Strikingly, we find that the ALS-associated FUS^P525L^ mutation profoundly disrupts the endosomal RNA landscape. Nearly all RNAs normally enriched at endosomes in wild-type neurons are lost in mutant cells. This collapse of the endosomal transcriptome coincides with reduced GEMIN5 association with endosomes. This, together with the finding that FUS interacts with GEMIN5, is consistent with a model in which the cytoplasmic mutant FUS sequesters GEMIN5 away from vesicular membranes. Moreover, the ability of mutant FUS to directly bind a substantial subset of endosomal RNAs suggests a dual mechanism of interference, involving both competition for RNA binding and disruption of GEMIN5-mediated targeting. These findings provide a molecular link between ALS-linked defects in RNA transport, organelle dynamics, and localized translation, and suggest that impaired endosomal RNA loading may contribute to synaptic and axonal dysfunction in the disease.

Together, our data establish a previously unrecognized mechanism for mRNA localization to late endosomes in neurons and reveal its vulnerability to ALS-associated perturbations. By defining the cis-acting features and trans-acting factors that govern RNA loading onto endosomal membranes, we provide a framework to understand how transcriptomic compartmentalization supports neuronal homeostasis.

## ACKNOWLEDGEMENTS

We thank M. Marchioni for technical help and M. Caruso for assistance. We thank Prof. M. Morlando and Prof. A. Rosa for discussion. This work was partially supported by grants from ERC-2019-SyG 855923-ASTRA, AIRC IG 2019 Id. 23053, “National Center for Gene Therapy and Drug based on RNA Technology” (CN00000041) and NextGenerationEU PNRR MUR to IB, together with PRIN 2022HM5LFW to JM. We thank Prof. Alberto Diaspro and Dr. Paolo Bianchini (Nanoscopy & NIC@IIT, Istituto Italiano di Tecnologia), Dr. Michele Oneto (Nikon Imaging Center) and Marco Scotto (Molecular Microscopy and Spectroscopy, Istituto Italiano di Tecnologia) for experimental support in confocal microscopy. We are also grateful to Dr. Diego Vozzi and the Genomic Facility of IIT for support in RNA sequencing experiments.

## AUTHOR CONTRIBUTIONS

I.B., D.M., A.S. and L.S.M. conceptualized the project. I.B., D.M., A.S. and J.M. supervised and coordinated the experiments. L.S.M. and D.M. set up the experimental system with the contribution of F.C. and performed molecular biology experiments. A.S. and A.D.A designed and performed the bioinformatic analyses. L.S.M., T.S., E.V. and D.M. performed the IF and smFISH experiments and designed the analysis. L.S.M., T.S. and V.P. analyzed the imaging experiments. J.M. analyzed the results and discussed the data. I.B., D.M., A.S. and L.S.M. wrote the original manuscript with contribution of all the authors.

## METHODS

### Cell culture, maintenance and treatments

SK-N-BE cells were cultured in adherence in RPMI-1640 medium supplemented with 10% fetal bovine serum (FBS), sodium pyruvate (1 mM), GlutaMAX™ (1X), penicillin (100 U/mL), and streptomycin (100 μg/mL) (all from Thermo Fisher Scientific). For long-term storage, cells were cryopreserved using a growth medium containing 20% FBS and 10% dimethyl sulfoxide (DMSO; Sigma-Aldrich) and rapidly recovered when required by thawing in a 37°C water bath. FUS overexpression was induced by incubating cells with 50 ng/mL doxycycline hyclate (Sigma-Aldrich) for 24 hours in culture medium. Induced pluripotent stem cells (iPSCs) were maintained as adherent cultures on dishes coated with Geltrex™ LDEV-free, hESC-qualified, reduced growth factor basement membrane matrix (1:100 dilution in serum-free medium; Gibco). Cells were cultured in NutriStem® hPSC XF medium (Sartorius) supplemented with penicillin (50 U/mL) and streptomycin (50 μg/mL) (Thermo Fisher Scientific). For cryopreservation, iPSCs were stored in NutriFreez® D10 Cryopreservation Medium and rapidly recovered by thawing in a 37°C water bath. Following thawing or enzymatic dissociation with Accutase (Thermo Fisher Scientific), cells were plated onto Geltrex™-coated dishes in complete culture medium supplemented with the ROCK inhibitor Y-27632 (10 μM; Enzo Life Sciences) for the first 24 h only to enhance cell survival. Colony morphology and growth were monitored daily by optic microscopy. For APEX2-mediated proximity labeling experiments, iPSC-derived motor neurons were incubated in neuronal maintenance medium consisting of Neurobasal medium supplemented with B27, brain-derived neurotrophic factor (BDNF, 20 ng/mL), glial cell line–derived neurotrophic factor (GDNF, 10 ng/mL; both from PeproTech), and L-ascorbic acid (200 ng/mL; Sigma-Aldrich). Cultures were preincubated with biotin-phenol (1 mM; IRIS Biotech) for 1 h at 37°C, followed by a brief exposure to hydrogen peroxide (1 mM H₂O₂ in PBS) for 1 min to activate APEX2 and initiate biotinylation. The reaction was immediately quenched by two washes with quenching buffer containing sodium ascorbate (10 mM) and Trolox (5 mM; Merck) in PBS.

### Cloning and stable cell lines generation

Isogenic iPSCs homozygous for either the FUS^WT^ or FUS^P525L^ alleles were kindly provided by our collaborator Prof. A. Rosa. Cell lines were generated as previously described^35^ and genetically engineered by random genomic integration of a cassette encoding the motor neuron transcription factors Ngn2-F2A-Isl1-T2A-Lhx3 (NIL) using an enhanced piggyBac transposable vector (epB-Bsd-TT-NIL). This system enables doxycycline-inducible differentiation into spinal motor neurons, as reported previously^23^. Plasmid encoding N-terminal FLAG-APEX2–tagged human RAB7A was obtained from Addgene (plasmid #135652). FLAG-APEX2-RAB7A insert was subcloned into the ePB-NEO-EIF1a backbone downstream of the constitutive EIF1a promoter. Insert was PCR-amplified using CloneAmp HiFi PCR Premix (TakaraBio) and cloned into acceptor vector linearized by inverse PCR using the InFusion Cloning system (TakaraBio), following the manufacturer’s guidelines for both reaction setup and primer design. To generate the GFP-tagged RAB7A, the FLAG-APEX2 coding sequence was excised from the ePB-NEO-EIF1a:FLAG-APEX2-RAB7A plasmid by inverse PCR and replaced with GFP amplified from a plasmid encoding N-terminal GFP-tagged human G3BP1, kindly provided by Roy Parker’s laboratory (University of Colorado Boulder). Cloning was performed using the InFusion Cloning system, yielding the ePB-NEO-EIF1a:GFP-RAB7A construct. For RLUC constructs, the Renilla luciferase (RLUC) coding sequence was amplified from the psiCHECK2 plasmid (Promega) and cloned into the inverse-PCR–linearized ePB-NEO-EIF1a backbone using the InFusion Cloning system, generating the ePB-NEO-EIF1a: RLUC plasmid. To insert the 5′ untranslated region of endosome-*enriched*, *depleted* or *unspecific* mRNAs upstream of the RLUC coding sequence, the corresponding 5′UTR was amplified from cDNA derived from WT motor neurons used in this study and cloned into the ePB-NEO-EIF1a:RLUC construct according to the manufacturer’s protocol, yielding ePB-NEO-EIF1a:5′UTR-RLUC plasmids. To establish stable iPSC lines, constructs were introduced via Lipofectamine™ Stem Reagent (Thermo Fisher Scientific), adhering to the transfection protocol recommended by the manufacturer. Successfully transfected cells were selected by neomycin resistance. SK-N-BE stable cell lines were generated as in Mariani et al.^19^

### iPSCs differentiation into spinal motor neurons

Human iPSCs were cultured under standard maintenance conditions and enzymatically dissociated into single cells using Accutase (Thermo Fisher Scientific). Cells were seeded onto Geltrex™-coated culture surfaces (1:100 dilution in serum-free medium; Gibco) at a density of 6.25 × 10⁴ cells/cm² in NutriStem® hPSC XF medium (Sartorius) supplemented with penicillin (50 U/mL), streptomycin (50 μg/mL), and the ROCK inhibitor Y-27632 (10 μM; Enzo Life Sciences). Twenty-four hours after plating, spinal motor neuron differentiation was initiated by doxycycline (1 μg/mL; Thermo Fisher Scientific)–mediated induction in DMEM/F12 (Sigma-Aldrich) containing penicillin, streptomycin, and GlutaMAX™ (Thermo Fisher Scientific). Following 48 h of induction, cultures were transitioned to Neurobasal/B27 differentiation medium consisting of Neurobasal Medium supplemented with B27, GlutaMAX™, non-essential amino acids (NEAA), penicillin, and streptomycin (all from Thermo Fisher Scientific). This medium was further supplemented with doxycycline (1 μg/mL), the γ-secretase inhibitor DAPT (5 μM; Sigma-Aldrich), and the FGFR inhibitor SU5402 (4 μM; Sigma-Aldrich). On day 5 of differentiation, cells were dissociated again with Accutase and replated onto Geltrex™-coated dishes or glass coverslips at a density of 1 × 10⁵ cells/cm². ROCK inhibitor (10 μM) was included for the first 24 h following replating to enhance cell survival. From day 5 onward, neuronal cultures were maintained in maturation medium composed of Neurobasal/B27 supplemented with brain-derived neurotrophic factor (BDNF, 20 ng/mL), glial cell line–derived neurotrophic factor (GDNF, 10 ng/mL; both from PeproTech), and L-ascorbic acid (200 ng/mL; Sigma-Aldrich). Cultures were maintained under these conditions until day 8, at which point cells were collected for downstream analyses.

### RNA extraction and analysis

Total RNA was isolated using the Direct-zol™ Miniprep RNA Purification Kit (Zymo Research), including an on-column DNase digestion step, according to the manufacturer’s instructions. RNA was eluted in 25 μL of RNase-free water. Reverse transcription was performed using the PrimeScript™ RT Reagent Kit (TakaraBio) following the supplier’s protocol. Quantitative PCR (qPCR) was carried out using PowerUp™ SYBR™ Green Master Mix (Thermo Fisher Scientific) in 15 μL reaction volumes on a QuantStudio™ 5 Real-Time PCR System (Thermo Fisher Scientific), employing standard cycling parameters. For GEMIN5-V5 CLIP experiments, data were reported as percentage of input. Briefly, Ct values obtained from input samples were corrected for sample dilution by subtracting the log₂ of the dilution factor. The percentage of input for each target transcript was subsequently calculated using the formula: % Input = 2^(Adjusted Input Ct−IP Ct).Primers used for qRT-PCR are listed in Table S3.

### Protein extraction and analysis

Protein extracts for quantification of total protein levels were obtained using RIPA buffer (50 mM Tris-HCl pH 7.5, 150 mM NaCl, 0.1% SDS, 0.5% sodium deoxycholate, 1% TRITON X-100, 5 mM EDTA), supplied with 1× Complete Protease Inhibitor Cocktail (Merck). For what concern protein extracts for APEX2-Pulldown experiments, RIPA buffer, supplied with 1× Complete Protease Inhibitor Cocktail (Merck), 1x Ribolock RNase Inhibitor (Thermo Fisher Scientific), 0.5 mM DTT, 10 mM sodium ascorbate (Merck) and 5 mM Trolox (Merck), was used. Protein concentration was assessed by Pierce™ 660 nm protein assay (Thermo Fisher Scientific), according to the manufacturer’s instructions. Protein electrophoresis was performed using 4–15% Mini-PROTEAN TGX Precast acrylamide gel (Bio-Rad), and proteins were transferred to 0,45 μm nitrocellulose membrane, using the TransBlot Turbo System (Bio-Rad), according to the manufacturer’s protocol. Membranes were stained with Ponceau, blocked with 5% non-fat dry milk (Sigma Aldrich) for 1 h and incubated overnight at 4°C with the following primary antibodies: anti-RAB7a (Abcam, AB137029, 1:1000), anti-LAMP1 (DHSB biology, H4A3, 1:1000), anti-GAPDH-HRP (Santa Cruz Biotechnology, sc-47724, 1:1000), anti-FLAG (Merck, F7425, 1:1000), anti-GEMIN5 (Sigma Aldrich, HPA037393, 1:500), anti-RAB5 (BD Transduction LaboratoriesTM, 610724, 1:1000), anti-H3 (Active motif, AB_2687567, 1C8B2, 1:1000), anti-EEA1 (eBioscienceTM, 14-9114, 1:1000), anti-TOM20 (Santa Cruz Biotechnology, sc-17764, 1:1000), anti-VINCULIN (ProteinTech, 26520-1-AP, 1:5000). After secondary antibody incubation for 1 hour at room temperature, protein detection was carried out either with Immobilon Crescendo Western HRP substrate (Merck), for total protein extract, or with Immobilon ECL UltraPlus Western HRP substrate, for APEX2-Pulldown experiments, using ChemiDoc™ MP System and images were analysed using Image Lab™ Software (Bio-Rad).

### APEX2-pulldown of proteins and RNAs

APEX2-mediated pulldown was performed with modifications to the protocol described by Kaewsapsak et al^21^. Cells expressing APEX2 were cultured and treated as described above; control samples lacking peroxidase activation were processed in parallel without hydrogen peroxide exposure. Following biotinylation, cells were washed twice and fixed with 0.1% paraformaldehyde (PFA) in quenching buffer for 10 min. Residual PFA was quenched by addition of glycine to a final concentration of 125 mM, followed by incubation with gentle agitation at room temperature for 5 min. Cells were subsequently washed twice and lysed directly on the culture dish in complete RIPA buffer. Cell lysates were incubated under rotation at 4°C for 15 min and then sonicated on ice at 20% amplitude for 30 s (1 s on/1 s off), followed by a 30 s pause; this cycle was repeated three times per sample. Debris was removed by centrifugation at 15,000 × g for 5 min at 4°C. Protein concentration was determined as described above, and 200 μg of total protein per biological replicate was used for pulldown experiments. Samples were adjusted to a final volume of 200 μL with complete RIPA buffer, and 5% of each sample was reserved as input. To each sample, 170 μL of NLB buffer (25 mM Tris-HCl pH 7.5, 150 mM KCl, 0.5% NP-40, 5 mM EDTA) supplemented with protease inhibitors (1× Complete Protease Inhibitor Cocktail, Merck), RNase inhibitor (1× Ribolock, Thermo Fisher Scientific), DTT (0.5 mM), and antioxidants (10 mM sodium ascorbate and 5 mM Trolox; Merck) were added. Streptavidin magnetic beads (40 μL; Pierce™, Thermo Fisher Scientific) were prewashed three times with binding and wash buffer (5 mM Tris-HCl pH 7.5, 1 M NaCl, 0.1% Tween-20, 0.5 mM EDTA), followed by two washes with a 1:1 mixture of RIPA and NLB buffers, and resuspended in the same buffer mixture. Beads were incubated with lysates for 2 h at 4°C under rotation (10 rpm). After binding, beads were washed eight times using buffers specified in the original protocol. For RNA-based downstream analyses, both pulldown and input samples were subjected to proteinase K digestion for 1 h at 42°C followed by an additional 1 h at 55°C. RNA was subsequently extracted using TRIzol™ LS (Thermo Fisher Scientific). For proteomic analyses, bound proteins were eluted after the final wash by incubation at 80°C for 10 min in 40 μL of elution buffer containing 1× Laemmli sample buffer (Bio-Rad), 50 mM DTT, and 2 mM free biotin (Merck).

### Library preparation and RNA-seq analysis

RNA libraries were prepared using the Illumina Stranded Total RNA Prep with Ribo-Zero Plus kit and sequenced on an Illumina NovaSeq 6000 platform in paired-end mode, generating reads of 100 bases. Read quality was assessed with FastQC v0.11.9, which revealed a transient reduction in quality at the first base, residual contamination consistent with incomplete ribosomal RNA removal by Ribo-Zero Plus, and the presence of Illumina adapter sequences. Adapter trimming and removal of low-quality nucleotides were performed using cutadapt v3.2^36^ with: *-u 1 -U 1 --trim-n --nextseq-trim=20 -m 35* and Trimmomatic software v0.39^37^ with PE mode and the following parameters; *ILLUMINACLIP:adapter_path:2:30:10:8:true LEADING:3 TRAILING:3 SLIDINGWINDOW:4:20 MINLEN:35*. To eliminate remaining ribosomal RNA, trimmed reads were aligned to an rRNA-only reference (**Table S1**) using Bowtie2 v2.4.2^38^, and only unmapped reads were retained. These filtered reads were subsequently aligned to the GRCh38 human genome using STAR^39^ v2.7.7a with the following parameters: *--outSAMstrandField intronMotif --outSAMattrIHstart 0 --outSAMtype BAM SortedByCoordinate --outFilterType BySJout --outFilterMultimapNmax 20 --alignSJoverhangMin 8 --alignSJDBoverhangMin 1 --outFilterMismatchNmax 999 --outFilterMismatchNoverLmax 0.04 --outFilterIntronMotifs RemoveNoncanonical --readFilesCommand zcat --outReadsUnmapped Fastx --peOverlapNbasesMin 35 --alignEndsType EndToEnd*. Because pull-down–derived RNA libraries typically undergo additional PCR amplification, PCR duplicates were removed using Picard MarkDuplicates v2.24.1(https://broadinstitute.github.io/picard/), and then gene level quantifications were performed with HTSeq-count^40^ v0.13.5 using Ensembl GTF gene annotation related to release 99^41^ and these parameters: *-s reverse -m union -t exon*. Read counts at each processing stage are summarized in the accompanying prospect (**Table S1**). Enrichment between biotin pull-down and inputINPUT samples was evaluated using DESeq2^42^ v1.34.0. Only transcripts with at least 1 average FPKM in input. To account for variability stemming from independent iPSC-to-motor neuron differentiation batches, each differentiation date was included as a batch factor in the DESeq2 design. Transcripts were classified as enriched when displaying a log₂ fold change greater than 0.59 with an adjusted P value below 0.05, and as depleted when the log₂ fold change was lower than –0.59 with an adjusted P value below 0.05. Transcripts not meeting these criteria were considered unspecific. This procedure was applied to both RAB7 H₂O₂–treated samples and the corresponding –H₂O₂ controls.

### RBP binding enrichment analysis

To identify RNA-binding proteins (RBPs) preferentially associated with the 5′UTRs of endosomal transcripts compared to depleted RNAs, we analyzed enhanced crosslinking and immunoprecipitation (eCLIP) datasets from K562 and HepG2 cells^27^. Of 150 profiled RBPs, those annotated as nuclear or with uncertain localization were excluded based on the Human Protein Atlas^43^ subcellular localization dataset (https://www.proteinatlas.org/humanproteome/subcellular/data#locations), yielding 49 cytoplasmic RBPs for downstream analyses. Endosome-enriched and depleted RNA sets were defined by enrichment analysis and restricted to protein-coding genes expressed in motoneurons, K562, and HepG2 cells. Genomic coordinates for 5′UTRs were extracted from Ensembl gene annotation release 99. Strand-specific intersections between RBP eCLIP peaks and 5′UTRs were performed using BEDTools v2.29.1^44^ to identify RBP–transcript interactions. For each RBP, transcripts with overlapping peaks on 5’UTR were classified as interacting, while transcripts lacking overlaps on their 5’UTR were considered non-interacting. Statistical significance of RBP binding enrichment to 5′UTRs of endosome-enriched versus depleted transcripts was assessed using Fisher’s exact tests. P values were corrected for multiple testing using false discovery rate (FDR) adjustment, and RBPs with FDR < 0.05 and log2OddsRatio > 1 were considered preferentially interacting with the 5’UTR of endosomal RNAs (**Table S2**). To assess 5′UTR specificity relative to coding sequence (CDS) and 3′UTR regions, we took advantage of metatranscript profile generated based on the lengths of the 5′UTR, CDS or 3′UTR regions of the expressed transcripts (FPKMs > 1). This information, together with the transcript sequences were retrieved from the Ensembl 99 database using the biomartR package v.2.50.3^45^. Each mRNA region was divided into an equal number of bins, ensuring the proportions among the lengths of the transcript’s regions were maintained. The bin ranges for each RNA were compiled into a BED file, and the sequences for each bin were extracted using BEDTools getfasta from the transcriptome FASTA file. For each RBP, their profiles were calculated based on the frequencies of interacting bins followed by ZScore normalization. Graphical representations of meta-region and meta-transcript analyses were performed using ComplexHeatmap R package v2.10.0^46^.

### Conservation Analysis

To assess evolutionary conservation of 5′UTRs in endosomal transcripts, we analyzed phylogenetic conservation scores across transcript regions. Longest isoform of protein-coding transcripts were selected, according to the ensemble annotation release 99. Transcript lengths were used to generate consecutive, non-overlapping 50-nt bins spanning full transcript bodies using BEDTools makewindow version 2.29.1. Transcript bins were annotated as 5′UTR, CDS, or 3′UTR and mapped to genomic coordinates in a strand-specific manner. Conservation scores were extracted for each bin using the BigWig file of phyloP100-way vertebrate conservation scores retrieved from UCSC^47^ golden path (https://hgdownload.cse.ucsc.edu/goldenpath/hg38/phyloP100way/), and mean phyloP scores were assigned per bin. Conservation scores were compared between 5′UTR and 3′UTR regions. Statistical differences in conservation between enriched and depleted transcript groups were assessed using the Mann–Whitney U test.

### Immunofluorescence analysis

Cells were fixed with 4% paraformaldehyde for 10 min at room temperature, followed by three washes with PBS. Membrane permeabilization was achieved by incubating coverslips in PBS containing 0.05% Triton X-100 for 5 min at room temperature. Non-specific binding was blocked by incubation in PBS supplemented with 2% bovine serum albumin (BSA) for 30 min at room temperature. Coverslips were then incubated for 1 h at RT with primary antibodies diluted in PBS containing 1% BSA. The following primary antibodies were used: anti-RAB7A (Abcam, ab137029; 1:250), anti-LAMP1 (Developmental Studies Hybridoma Bank, H4A3; 1:500), anti-FLAG (Merck, F3165; 1:500), anti-FLAG (Merck, F3165, 1:500) anti-GEMIN5 (Sigma-Aldrich, HPA037393; 1:250), anti-DCP1a (Santa Cruz Biotechnology, sc-100706, 1:250). After three washes in PBS (5 min each), samples were incubated for 45 min at room temperature with fluorophore-conjugated secondary antibodies (Thermo Fisher Scientific) diluted 1:300 in PBS containing 1% goat or donkey serum. Following secondary antibody incubation, coverslips were washed three times with PBS and nuclei were counterstained with DAPI (1 μg/mL in PBS; Sigma-Aldrich) for 3 min at room temperature. Coverslips were mounted onto microscope slides using ProLong™ Diamond Antifade Mountant (Thermo Fisher Scientific, P36961). Imaging was performed using a Nikon A1 confocal laser-scanning microscope equipped with 60× and 100× objectives. Z-stack images were acquired using the NIS-Elements AR software (Nikon) with the ND Acquisition module, using a Z-step size of 150–175 nm. Image analysis was conducted using FIJI/ImageJ open-source software. For colocalization analysis, nuclear regions were segmented based on the DAPI signal and excluded to restrict analyses to the cytoplasmic compartment. After Gaussian blur filtering and background subtraction, Pearson’s correlation coefficient was calculated on Maximum Intensity Projections using the JaCoP plugin. For particle count analysis, after Gaussian blur filtering and background removal, the analyse particles function was used and SOMA vesicles were identified by enlarging the mask of nuclei on the Maximum Intensity Projections of Z-planes. 3D details were produced using the 3D volume viewer plugin of Fiji software.

### Single-molecule fluorescence in situ hybridization analysis

RNA-FISH staining was carried out using the HCR RNA-FISH technology^48^ according to manufacturer protocol (https://www.molecularinstruments.com/hcr-rnafish), with minor modifications to improve simultaneous RNA and protein visualization. HCR probes targeted to the RNAs of interest and compatible with the B3 amplifier were designed using the Özpolat Lab HCR probe generators^49^. Probe sequences are available in Table S3. Imaging was performed as specified above. FIJI/ImageJ open-source software was used to analyse images. Colocalization analyses were carried out excluding nuclear regions as previously indicated. Gaussian blur filtering and background subtraction were applied before merging all Z-stacks with maximum intensity projection. RS-FISH plugin^50^ was used for RNA FISH spot detection. The % of FISH spot colocalization was calculated with the JaCoP plug-in tool. Digital enlargements were obtained by manual cropping from original full-field images. 3D details were produced using the 3D volume viewer plugin of Fiji software.

### UV-crosslinked immunoprecipitation (CLIP)

5×10^6^ SK-N-BE cells were seeded in a 15 cm plate. Cells were cultured for two days and UV-crosslinked on ice at 150 mJ/cm^2^, then collected by scraping in PBS and flash-frozen in dry ice. Pellets were lysed in CLIP Lysis Buffer (20 mM TRIS-HCl pH 7.5, 150 mM NaCl, 5 mM MgCl_2_, 0,5% NP-40, 0,5 mM DTT, 1x Protease Inhibitory Cocktail and 1:300 Ribolock (Thermo Fisher Scientific) by incubation for 10 minutes on ice, and lysates were homogenized through sonication at 25% amplitude for 5 pulses (2 s on/2 s off), followed by a 30 s pause; this cycle was repeated four times per sample. Lysates were cleared by centrifugation at 15000g, 10 minutes, 4°C. After BCA quantification, 500 µg of lysate were combined with 2 µg of anti-V5 antibody (Novus Biologicals, NB600-381) or control IgGs (Thermo Fisher, 02-6102), and incubated for 2:30 hours at 4°C in a final volume of 500 µL. Antibody-protein complexes were recovered by adding 50 µL of Dynabeads Protein G magnetic beads (Thermo Fisher Scientific) and incubating for 1 hour at 4°C. Beads were washed thrice in 1 mL of Wash Buffer (20 mM TRIS-HCl pH 7.5, 250 mM NaCl, 1mM MgCl_2_, 0,5% NP-40). ¾ of beads were treated with 100 µg of Proteinase K (Thermo Fisher Scientific, AM2546) in PK Buffer (75 mM TRIS-HCl pH 7.5, 10 mM EDTA, 0,1% SDS) for 1 hour at 42°C; the reaction was stopped by adding 5 volumes of TRIZOL LS Reagent (Thermo Fisher Scientific). The remaining ¼ of beads were used for protein elution by boiling in 1x Laemmli Sample Buffer, 50 mM DTT for 15 minutes at 90°C.

### Co-immunoprecipitation

5×10^6^ SK-N-BE cells were seeded in two 15 cm plates per sample. Cells were cultured for two days, collected by scraping in PBS and flash-frozen in dry ice. Pellets were lysed in 400 µL of Co-IP Buffer (1x PBS, 0.01% NP-40, 1x Protease Inhibitory Cocktail) on ice for 10 minutes, and lysates were homogenized by passing 10 times through a 26G needle. Lysates were cleared by two rounds of centrifugation at 5000g, 10 minutes, 4°C to obtain a cytoplasm-enriched fraction. After BCA quantification, 1 mg of lysate was combined with 50 µL of Dynabeads Protein G magnetic beads (Thermo Fisher Scientific) previously coupled with 2 µg of anti-V5 antibody (Novus Biologicals, NB600-381) or control IgGs (Thermo Fisher, 02-6102), and incubated for 4 hours at 4°C in a final volume of 500 µL. Beads were washed thrice with 1 mL of Co-IP buffer in rotation for 5 minutes at 4°C, and proteins were eluted by boiling in 1x Laemmli Sample Buffer, 50 mM DTT for 15 minutes at 90°C.

### Nucleo-cytoplasmic fractionation

Nucleo-cytoplasmic fractionation was carried out as specified in Vitiello et al^51^.

