## Supplementary Figures and Legends for "Neuronal late endosomes serve as selective RNA hubs disrupted by ALS-linked FUS mutation"

### Suppl. Figure 1

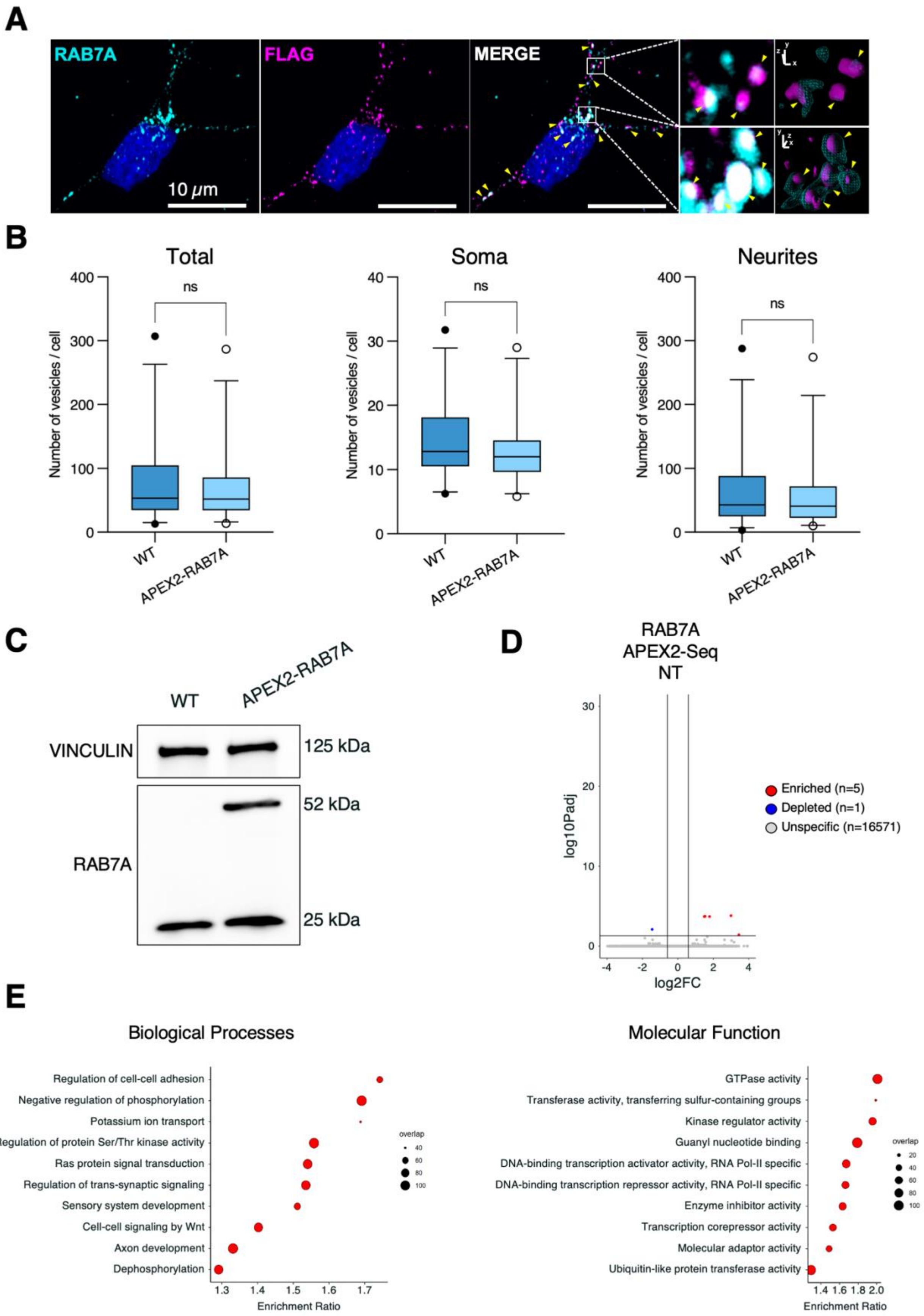

**F**

#### Endocytosis

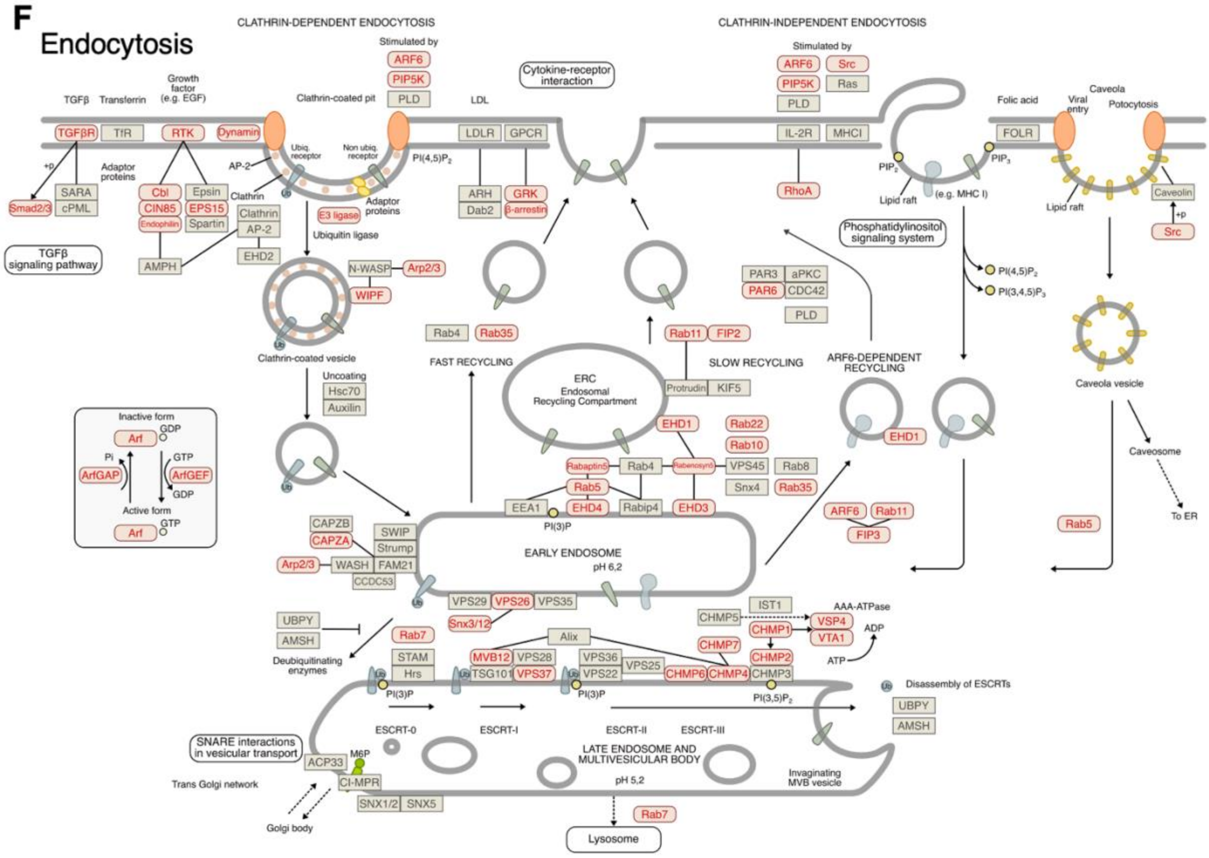


**G**

#### Axon guidance

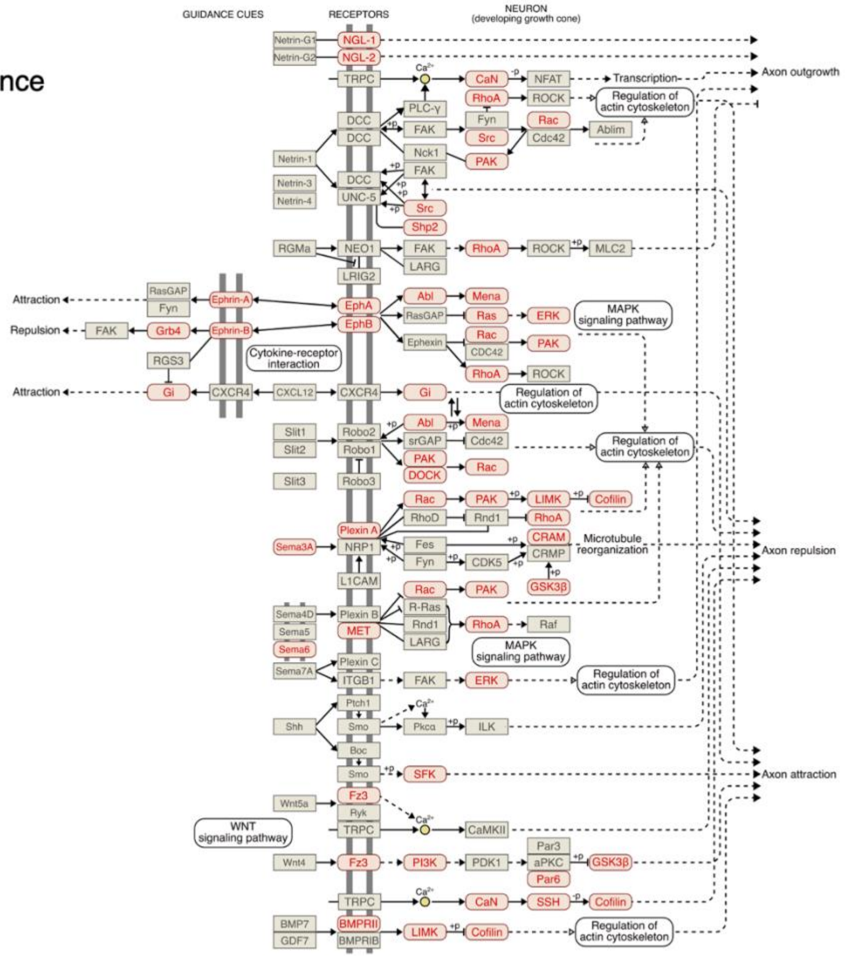

H

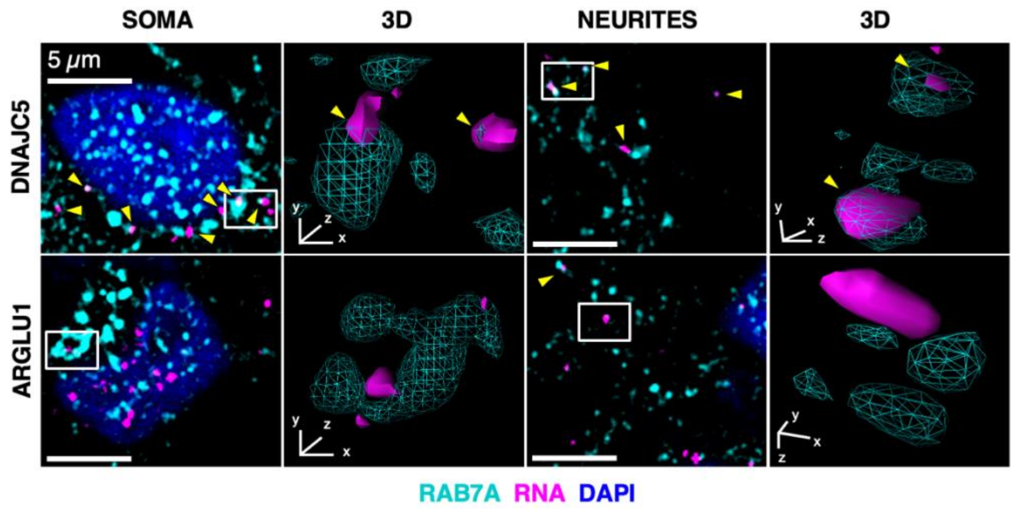
SOMA
3D
NEURITES
3D
5 μm
DNAJC5
ARGLU1
RAB7A RNA DAPI

I

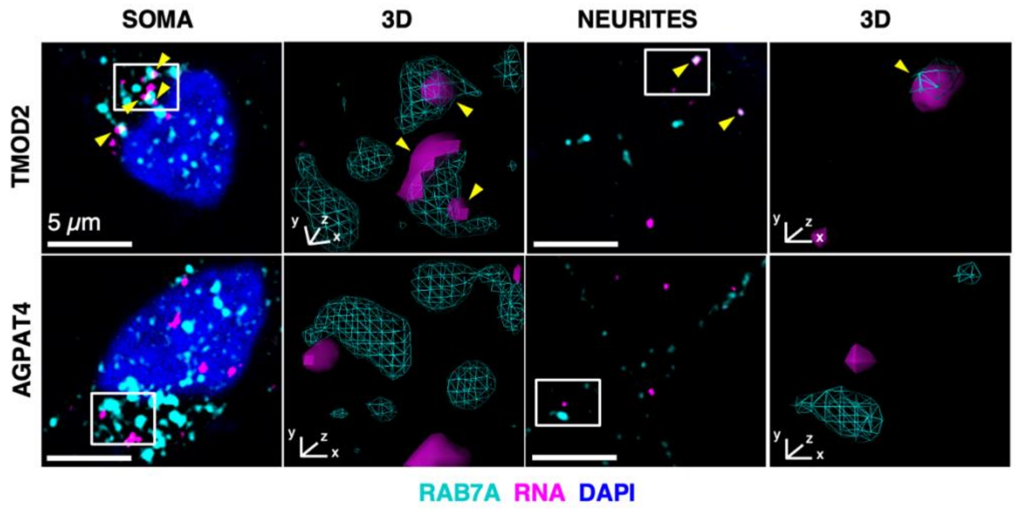
SOMA
3D
NEURITES
3D
5 μm
TMOD2
AGPAT4
RAB7A RNA DAPI

**Supplementary Figure 1. (A)** Immunofluorescence showing APEX2-RAB7A (FLAG, magenta) relative to endogenous RAB7A (cyan). Scale bar = 10 µm. 2D and 3D Details of selected boxed regions are shown. Colocalization is indicated by yellow arrows. **(B)** Quantification of total, somatic, and neuritic RAB7A-particles number per cell in wild-type and stable APEX2-RAB7A expressing MNs. Data are shown as boxplots. Statistical significance was assessed with two-tailed, unpaired T-test. N = 6 biological replicates (total cells: WT MNs n = 268, APEX2-RAB7A MNs n = 262) **(C)** Western blot showing relative APEX2-RAB7A and endogenous RAB7A protein levels in iPSCs. Vinculin serves as loading control. **(D)** Volcano plot showing the log2FC pull-down/input and the −log10Pvalue for each RNA detected in APEX2-RAB7A untreated (NT) condition. Enriched and Depleted RNAs (Padj < 0.05, |log2FC| > 0.59) or Unspecific RNAs are indicated by red, blue and gray dots, respectively. **(E)** Gene Ontology (GO) Biological Processes and Molecular Function analyses of RNAs enriched in the APEX2–RAB7A + $H_2O_2$ condition. **(F)** KEGG pathway annotation of endosome-enriched RNAs involved in endocytosis (left) and axonal development (right); pathway-associated genes are highlighted in red**. (G)** Additional representative RNA FISH and immunofluorescence images showing localization of endosome-*enriched* (DNAJC5) or endosome-*depleted* (ARGLU1) mRNA (magenta) relative to RAB7A-positive endosomes (cyan) in soma and neurites. Insets show 3D renderings of boxed regions. Scale bar = 5 µm. Colocalization is indicated by yellow arrows. **(H)** Representative RNA FISH and immunofluorescence images showing localization of endosome-*enriched* (TMOD2) or endosome-*unspecific* (AGPAT4) mRNA (magenta) relative to RAB7A-positive endosomes (cyan) in soma and neurites. Insets show 3D renderings of boxed regions. Scale bar = 5 µm. Colocalization is indicated by yellow arrows.

### Suppl Fig 2

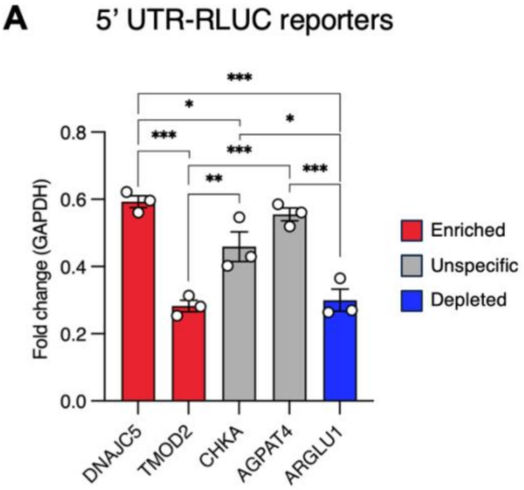
A
5' UTR-RLUC reporters
***
*
*
***
***
***
**
***
0.8
0.6
0.4
0.2
0.0
Fold change (GAPDH)
DNAJC5
TMOD2
CHKA
AGPAT4
ARGLU1
Enriched
Unspecific
Depleted

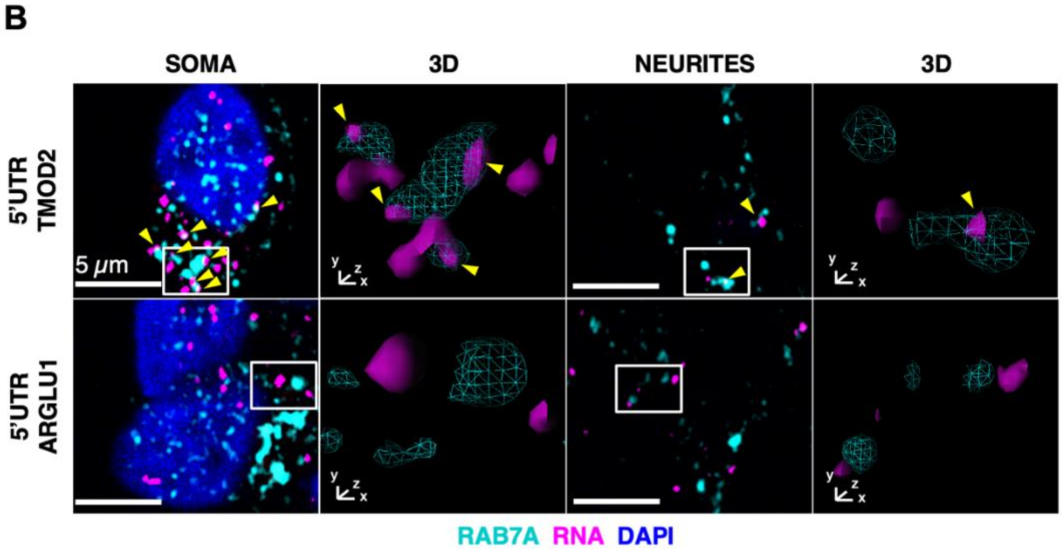
B
SOMA
3D
NEURITES
3D
5'UTR TMOD2
5'UTR ARGLU1
5 μm
RAB7A RNA DAPI

**Supplementary Figure 2. (A)** qPCR analysis of RLUC reporter mRNA expression levels in iPSC-derived MNs; GAPDH mRNA serves as a reference. Data are shown as mean ± SEM (N = 3). Statistical significance was assessed with ordinary two-way ANOVA test. **(B)** Representative RNA FISH and immunofluorescence images showing localization of RLUC reporter mRNAs (magenta) fused to an endosome-*enriched* (TMOD2) or *depleted* (ARGLU1) 5′UTR relative to RAB7A-positive endosomes (cyan) in soma and neurites. 3D rendering of selected white boxes showing RAB7A-particles (cyan) and reporter mRNA (magenta). Scale bar = 5 µm. Colocalization is indicated by yellow arrows.

#### Suppl fig.3

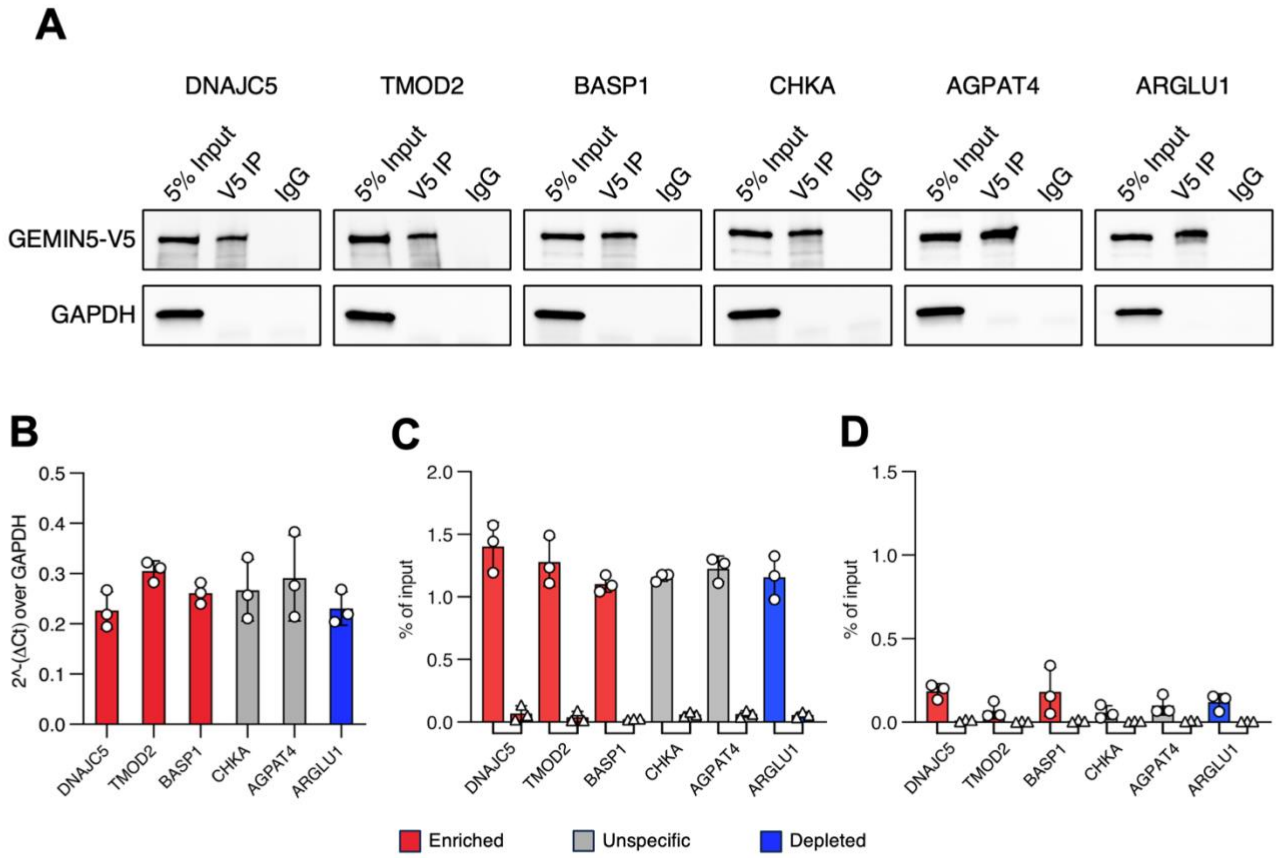
A
DNAJC5
TMOD2
BASP1
CHKA
AGPAT4
ARGLU1
5% Input
V5 IP
IgG
GEMIN5-V5
GAPDH
B
2^-(ΔCt) over GAPDH
C
% of input
D
% of input
DNAJC5
TMOD2
BASP1
CHKA
AGPAT4
ARGLU1
Enriched
Unspecific
Depleted

E

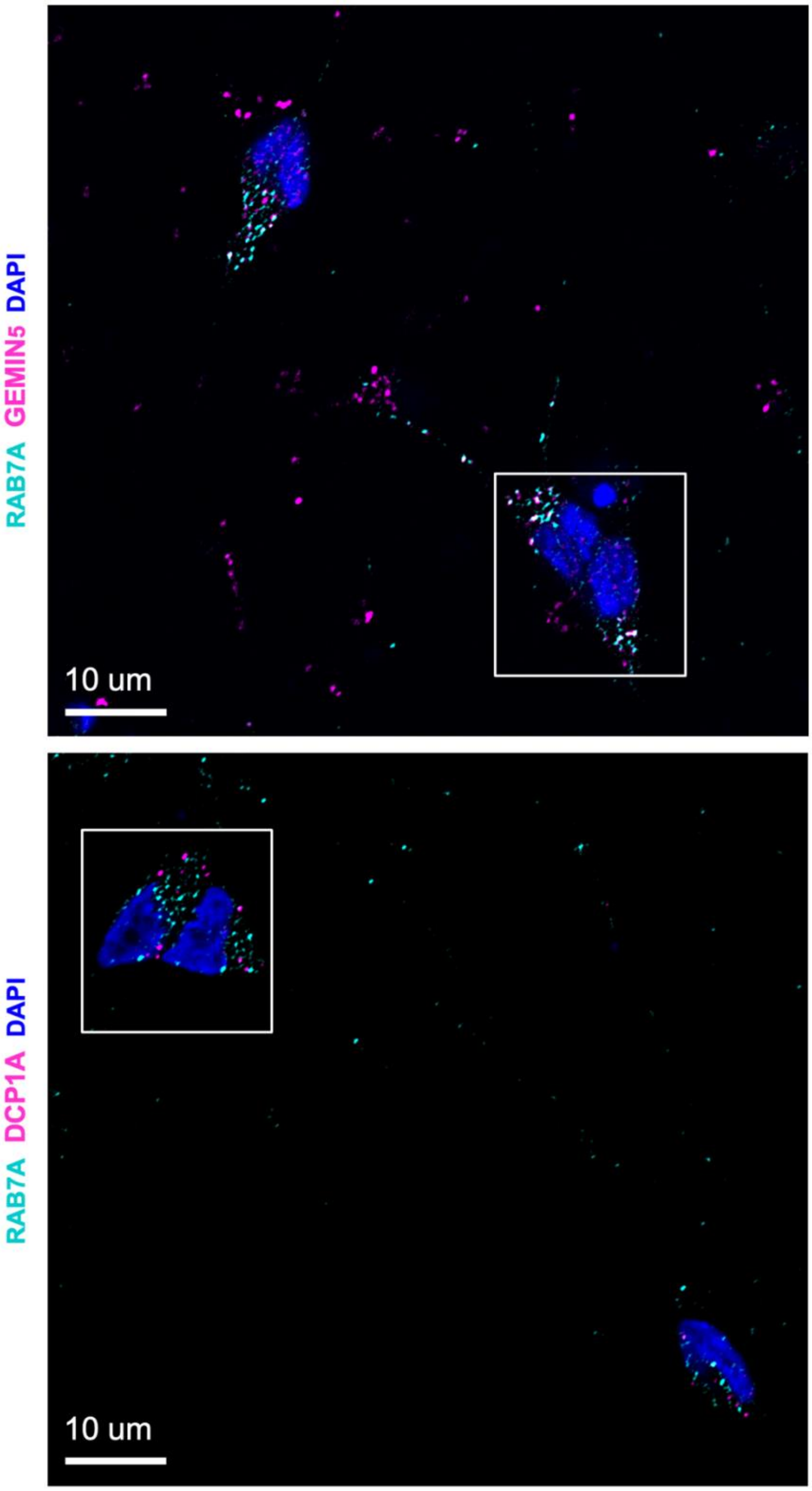

**Supplementary Figure 3. (A)** Representative western blot analysis of V5-GEMIN5 IP from neuroblastoma cell lines expressing different RLUC reporters. Immunoprecipitation (V5-IP) and control (IgG) samples are compared to 5% total protein Input. GAPDH serves as a negative control. **(B)** qPCR analysis of RLUC reporter mRNA expression levels in SK-N-BE cells; GAPDH mRNA serves as a reference. Data are shown as mean ± SEM (N = 3). **(C)** qPCR quantification of V5–GEMIN5 binding to endogenous GEMIN5 mRNA, used as a positive control, demonstrating comparable immunoprecipitation efficiency across reporter cell lines. Data are expressed as percentage of input. Error bars represent ± SEM (N = 3). **(D)** qPCR quantification of V5-GEMIN5 binding to GAPDH mRNA, serving as negative control, across reporter cell lines. Data is expressed as percentage of input. Error bars represent ± SEM (N = 3). **(G)** Uncropped selected slice of immunofluorescence image corresponding to Figure 3F, GEMIN5 (upper panel, magenta) or DCP1A (lower panel, magenta) relative to RAB7A-positive endosomes (cyan).

**Suppl. Fig. 4**

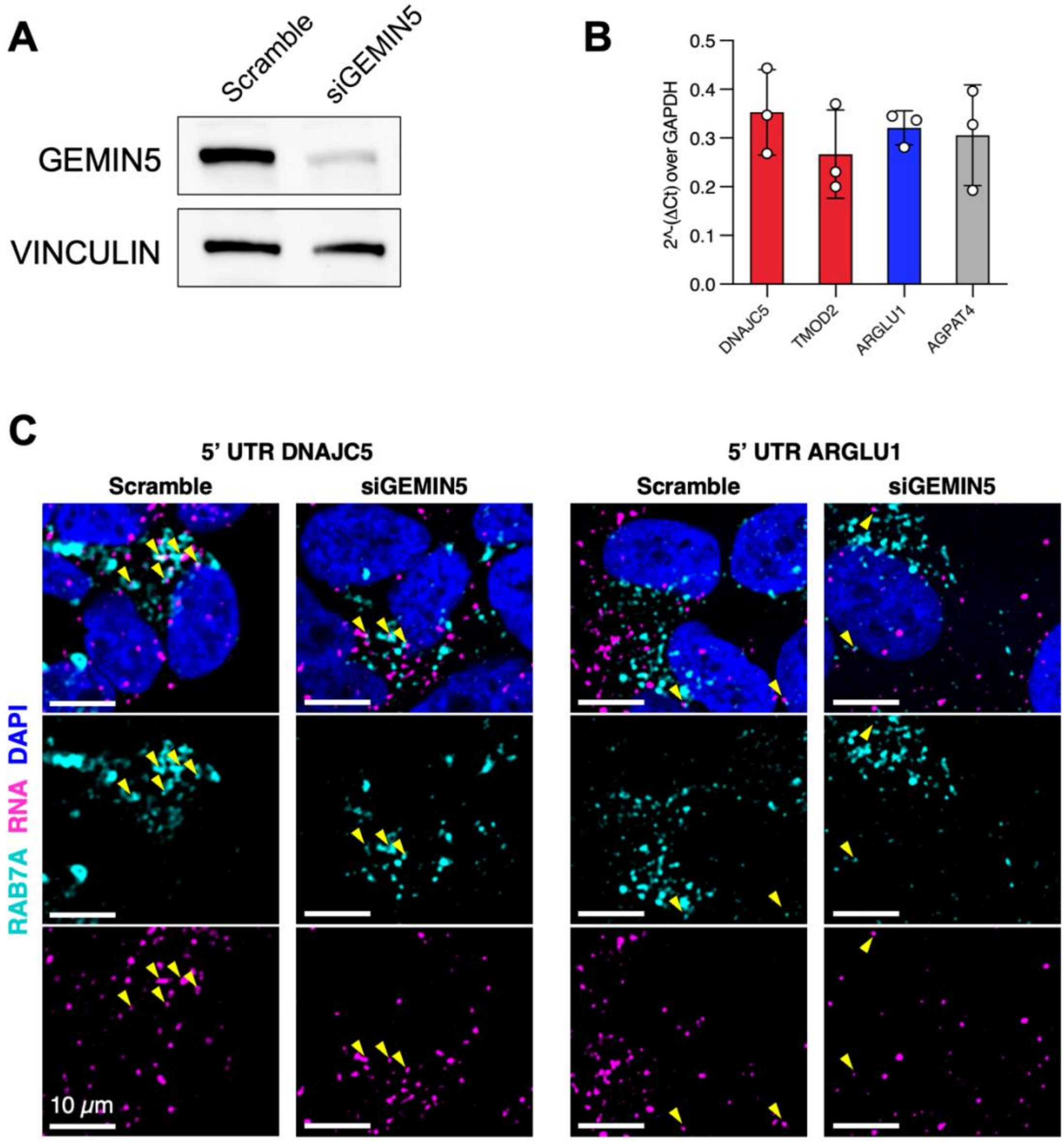
A
Scramble
siGEMIN5
GEMIN5
VINCULIN
B
0.5
0.4
0.3
0.2
0.1
0.0
2^-(ΔCt) over GAPDH
DNAJC5
TMOD2
ARGLU1
AGPAT4
C
5' UTR DNAJC5
5' UTR ARGLU1
Scramble
siGEMIN5
Scramble
siGEMIN5
RAB7A RNA DAPI
10 μm

**Supplementary Figure 4. (A)** Western blot showing GEMIN5 KD efficiency in SK-N-BE cells. Vinculin serves as loading control. **(B)** qPCR analysis of RLUC reporter mRNA expression levels in control and GEMIN5 KD SK-N-BE cells; GAPDH mRNA serves as a reference. Data are shown as mean ± SEM (N = 3). **(C)** Representative RNA FISH and immunofluorescence images showing localization of RLUC reporter mRNAs (magenta) fused to an endosome-*enriched* (DNAJC5) or *depleted* (ARGLU1) 5′UTR relative to RAB7A-positive endosomes (cyan) in control (Scramble) and GEMIN5 knockdown (siGEMIN5) SK-N-BE cells. Scale bar = 10 µm. Colocalization is indicated by yellow arrows.

**Suppl Fig.5**

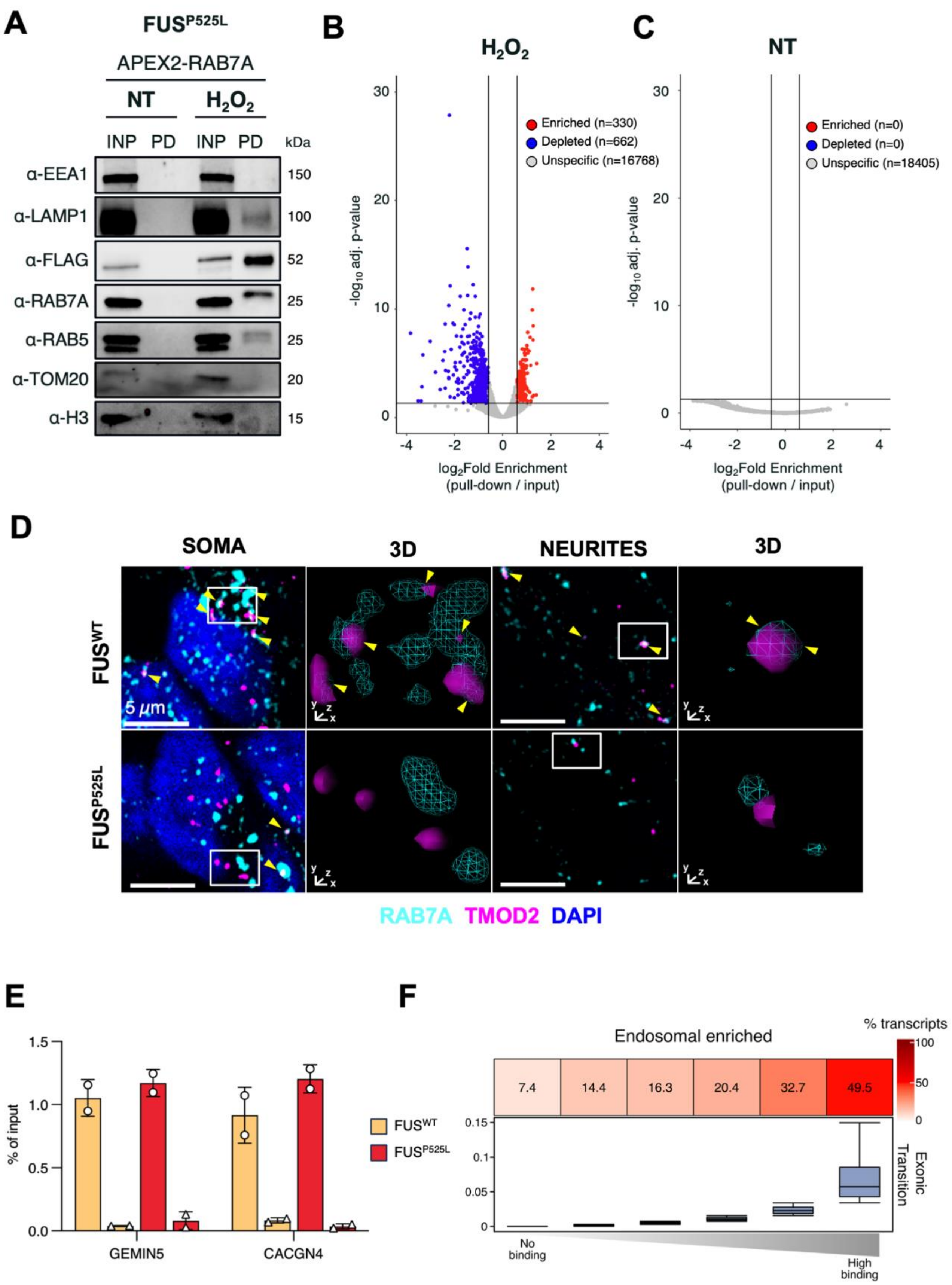

A
FUSP525L
APEX2-RAB7A
NT
H2O2
INP PD INP PD kDa
α-EEA1 150
α-LAMP1 100
α-FLAG 52
α-RAB7A 25
α-RAB5 25
α-TOM20 20
α-H3 15
B
H2O2
Enriched (n=330)
Depleted (n=662)
Unspecific (n=16768)
-log10 adj. p-value
log2Fold Enrichment (pull-down / input)
C
NT
Enriched (n=0)
Depleted (n=0)
Unspecific (n=18405)
D
SOMA
3D
NEURITES
FUSWT
FUSP525L
5 µm
RAB7A TMOD2 DAPI
E
% of input
FUSWT
FUSP525L
GEMIN5
CACGN4
F
Endosomal enriched
% transcripts
7.4 14.4 16.3 20.4 32.7 49.5
Exonic Transition
No binding
High binding

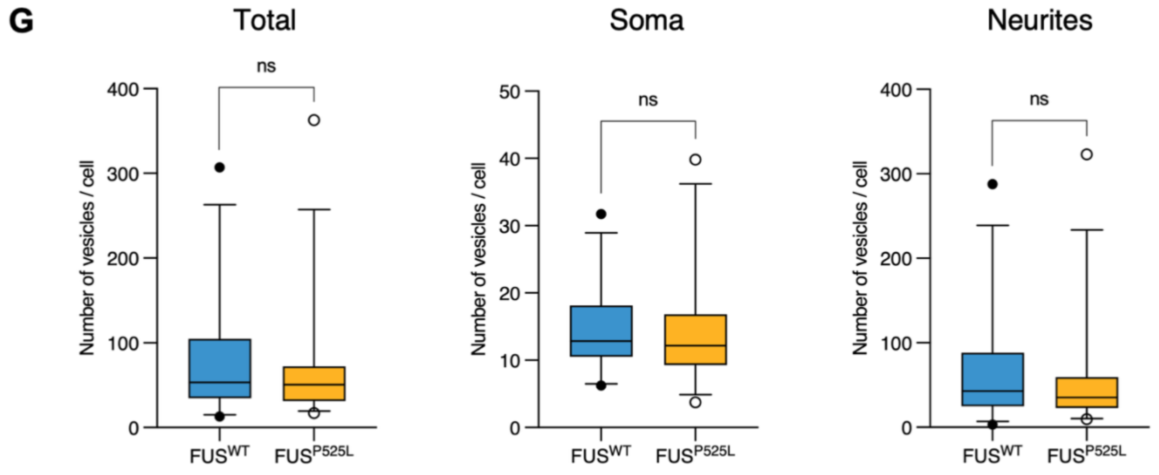
G
Total
Soma
Neurites
ns
ns
ns
Number of vesicles / cell
Number of vesicles / cell
Number of vesicles / cell
400
300
200
100
0
50
40
30
20
10
0
400
300
200
100
0
FUS^WT
FUS^P525L
FUS^WT
FUS^P525L
FUS^WT
FUS^P525L

**Supplementary Figure 5. (A)** Western blot analysis of the APEX2-pulldown on $FUS^{P525L}$ iPSCs-derived MNs expressing flagged APEX2-RAB7A protein. Early endosomal markers (EEA1, RAB5), Late endosomal marker (RAB7A), Late endosomal/lysosomal marker (LAMP1), APEX2-RAB7A construct (FLAG), Nuclear marker (H3), and mitochondrial marker (TOM20) were used to assess compartment purification. INP = 2,5% input, PD = pull down, H2O2= hydrogen peroxide treatment. **(B)** Volcano plot showing the log2FC pull-down/input and the −log10Pvalue for each RNA detected in FUSP525L APEX2-RAB7A + H2O2 condition. Enriched and Depleted RNAs (Padj < 0.05, |log2FC| > 0.59) or Unspecific RNAs are indicated by red, blue and gray dots, respectively. **(C)** Volcano plot showing the log2FC pull-down/input and the −log10Pvalue for each RNA detected in FUSP525L APEX2-RAB7A untreated (NT) condition. Enriched and Depleted RNAs (Padj < 0.05, |log2FC| > 0.59) or Unspecific RNAs are indicated by red, blue and gray dots, respectively. **(D)** Representative RNA FISH and immunofluorescence images showing TMOD2 mRNA (magenta) relative to RAB7A-positive endosomes (cyan) in soma and neurites, in $FUS^{WT}$ and $FUS^{P525L}$ MNs. Scale bar = 5 μm. Colocalization is indicated by yellow arrows. 3D rendering of selected white boxes showing RAB7A-particles (cyan) and TMOD2 mRNA (magenta). **(E)** qPCR quantification of GEMIN5 binding to CACNG4 mRNA in $FUS^{WT}$ and $FUS^{P525L}$ SK-N-BE cell lines, expressed as percentage of input. Data are shown as mean ± SEM (N = 2). **(F)** Transcripts were binned into six categories according to increasing $FUS^{P525L}$-binding signal: *no binding (n*=5183*)*, *low (n*=1643*), medium-low (n*=1643*)*, *medium (n*=1642*)*, *medium-high (n*=1642*) and high (n*=1643*)*. The top panel shows, for each bin, the percentages of transcripts classified as endosomal-enriched (colour intensity reflects % transcripts). The bottom panel displays the corresponding distribution of the exonic PAR-CLIP transition score across bins **(G)** Quantification of total, somatic, and neuritic RAB7A-particles number per cell in $FUS^{WT}$ and $FUS^{P525L}$ MNs. Data are shown as boxplots. Statistical significance was assessed with two-tailed, unpaired T-test. N = 6 biological replicates (total cells: $FUS^{WT}$ MNs n = 268, $FUS^{P525L}$ MNs n = 294)
